# Uncovering the Dynamics of Precise Repair at CRISPR/Cas9-induced Double-Strand Breaks

**DOI:** 10.1101/2023.01.10.523377

**Authors:** Daniela Ben-Tov, Fabrizio Mafessoni, Amit Cucuy, Arik Honig, Cathy Melamed-Bessudo, Avraham A. Levy

**Author notes:** These authors contributed equally to this work (Daniela Ben-Tov, Fabrizio Mafessoni).

## Abstract

CRISPR/Cas9-mediated genome editing relies on error-prone repair of targeted DNA double-strand breaks (DSBs). Understanding CRISPR/Cas9-mediated DSB induction and subsequent repair dynamics requires measuring the rate of cutting and that of precise repair, a hidden-variable of the repair machinery. Here, we present a molecular and computational toolkit for multiplexed quantification of DSB intermediates and repair-products by single-molecule sequencing. Using this approach, we characterized the dynamics of DSB induction, processing and repair at endogenous loci along a 72-hour time-course in tomato protoplasts. Combining this data with kinetic modeling reveals that indel accumulation is not an accurate reflection of DSB induction efficiency due to prominent precise re-ligation, accounting for 40-70% of all repair events. Altogether, this system exposes previously unseen flux in the DSB repair process, decoupling induction and repair dynamics, and suggesting an essential role of high-fidelity repair in limiting CRISPR editing efficiency in somatic cells.

## Introduction

DNA DSBs are one of the most cytotoxic forms of DNA damage, both endangering genome stability and driving genome evolution, through error-prone repair. In recent years, the ability to target DSBs to specific endogenous sequences, using custom-designed nucleases, in particular, with Clustered Regulatory Interspaced Short Palindromic Repeat associated protein Cas9 (CRISPR-Cas9) has triggered a revolution in genome engineering^1–5^, widely impacting the field of life science from biomedical technology and research^6–10^ to agriculture^11–15^.

Optimization of this technology for precise genome engineering requires understanding DSB induction and the dynamics of repair by the endogenous machinery. Once a DSB is made, it is repaired by a complex network of highly-conserved pathways, which can be characterized by the scars they leave behind (indels or conversion tracts resulting from templated repair)^16^. These pathways, generally subdivided into non-homologous end-joining (NHEJ) and homologous recombination (HR), have been extensively studied in organisms from yeast to plants and mammals^16–23^. While canonical NHEJ (cNHEJ) is the primary pathway in somatic plant and mammalian cells, other NHEJ pathways, can act_at a DSB site, involving microhomologies and/or DNA synthesis, when direct re-ligation through the canonical pathway has either failed, or is unavailable ^24–26^. These pathways have been referred to as alternative-EJ (alt-EJ), microhomology-mediated end-joining (MMEJ), or Polymerase Theta mediated End-Joining (TMEJ). In addition, our lab and others have shown that HR can act in somatic cells to repair DSBs through inter-homolog recombination^27–29^, or can lead to chromosomal rearrangement.^30,31^

Since DNA is constantly exposed to damage ^32^, maintaining genome integrity and limiting rapid accumulation of somatic mutations would require high-fidelity repair^33^. Accumulating evidence supports the suggestion that NHEJ may be highly accurate in mammalian systems ^34–38^. For example, DSBs induced by *I-SceI* were shown to be repaired precisely up to ~75% of the time in mouse cells^38^. In contrast, work characterizing re-ligation of linearized plasmids in both tobacco explants and in protoplasts suggests that NHEJ is highly error-prone, often displaying evidence of ‘filler DNA’^39^. Direct comparison of repair-fidelity in plants and mammals supports these results, measuring 50-55% precise repair in HeLa cells, compared to only 15-30% in tobacco cells^40^. However, these studies were largely conducted in the context of exogenous or transgenic DNA.

Studies of DSBs induced by CRISPR/Cas9 in endogenous chromatin have primarily focused on mutagenic repair, revealing many aspects affecting editing outcome and repair-pathway choice^41,42^. In addition, kinetic studies have provided insight into the dynamics of the process of induction and error-prone repair ^43–46^. However, as a result of the inability to directly observe scar-less re-ligation, measuring the degree of precise repair, particularly by the end-joining pathways, remains challenging. In addition, few studies follow the process of DSB induction and the intermediates between break and repair, namely the processing of the DSBs that leads to specific types of outcomes. Despite efforts, much of the process of CRISPR/Cas9-induced DSB repair remains unknown, particularly in somatic plant cells, including the efficiency and timing of induction, characteristics of DSB intermediates that lead to the different outcomes, and the fidelity of repair.

Resolving precise repair at CRISPR/Cas9 induced DSBs, in endogenous chromatin, requires quantification of both intermediates and products along a timecourse, facilitating estimation of the rates of cutting and repair, both precise and error-prone, through mathematical modeling. Using measurements of repair-error, quantified through amplicon sequencing, and DSB formation through ligation-mediated PCR, one study estimated cutting and repair rates in four CRISPR-Cas9 targets in K562 cell lines ^45^. Their results suggest that repair is often slow and highly error-prone, with an estimated half-life of up to ~10 hours ^45^. However, this approach is limited by resolution achievable in quantifying both DSBs and repair products, leading to estimates of precise repair accompanied by a large degree of uncertainty^45^. Reducing the uncertainty in the rate constant estimations requires improving the precision and resolution of measuring both DSBs and products of repair error.

Here, we present UMI-DSBseq, a novel molecular and computational toolkit for direct quantification of DSB intermediates alongside repair-products through multiplexed single-molecule sequencing. Using this approach, we follow the dynamics of DSB induction, end-processing, and repair at three CRISPR/Cas9 targets in tomato protoplasts, measuring the abundance of DSB intermediates and repair products, simultaneously, along a 72-hour time-course. In addition to revealing characteristics of DSB intermediates, including evidence for local off-target cleavage, combining this data with kinetic modeling reveals precise re-ligation as a prominent feature at all 3 targets, accounting for 40-70% of all repair events. Finally, we follow the processing of unrepaired DSBs, revealing the contribution of processed intermediates to error-prone repair. Altogether, these results suggest that editing efficiency in plants is determined, in some cases as much by the fidelity of the endogenous repair process as by the ability to efficiently induce DSBs at the target site.

## Results

### Single-molecule quantification and characterization of DSB intermediates and error-prone repair products using UMI-DSBseq

Evaluating the process of DSB repair, from intact to final repair-outcome, requires controlled, relatively synchronized DSB induction, followed by quantitative characterization of both DSB-repair intermediates and repair-products. To this end, preassembled ribonucleoproteins (RNPs), consisting of SpCas9 protein and synthetic sgRNA, are delivered directly into plant protoplasts purified from tomato seedlings using PEG-mediated transformation (Fig. 1A). This method, which does not rely on transcription, translation by the endogenous machinery, or assembly of Cas9 with the sgRNA *in vivo*, bypasses the lag-time between introduction and activity, allowing for fast induction of DSBs in a controlled manner. Following RNP delivery, we can follow dynamics of DSB induction and repair process along a time-course.

**Figure 1.**
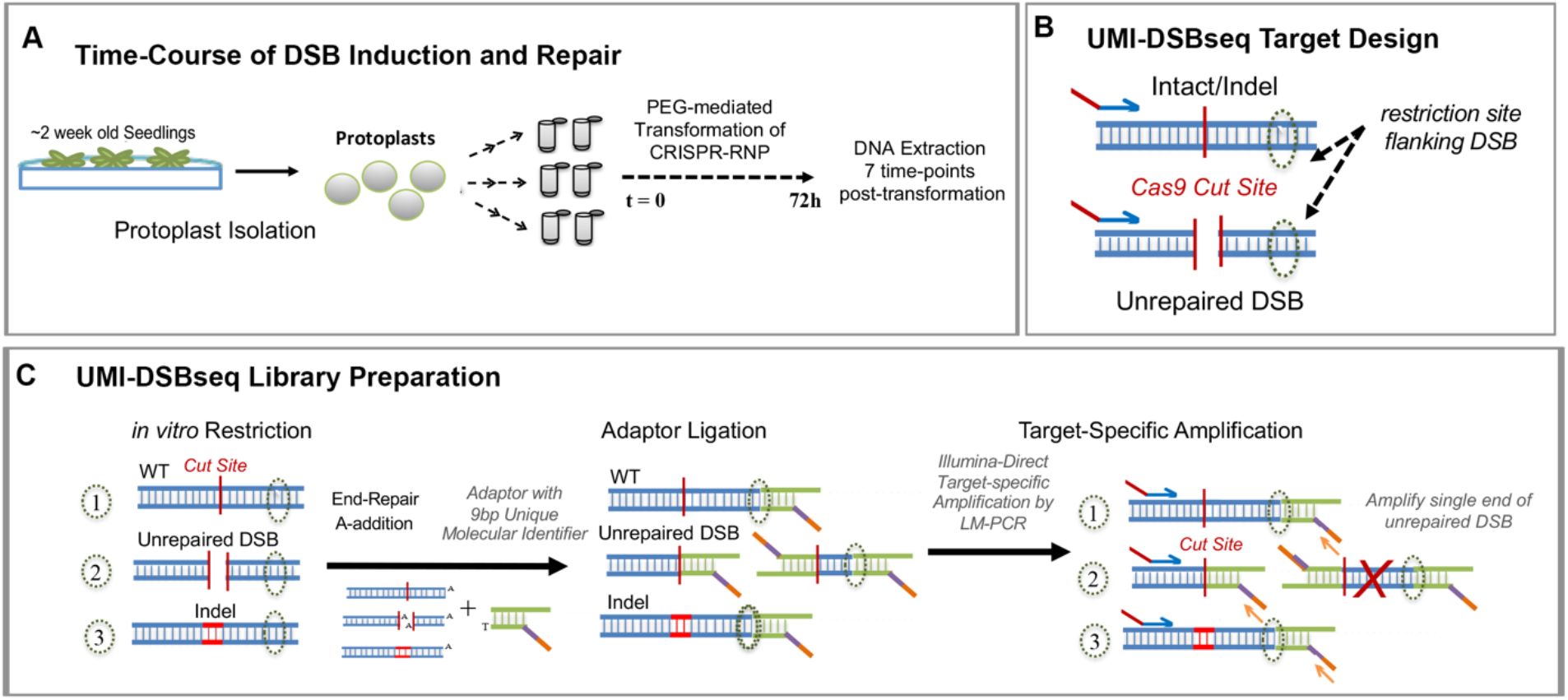
UMI-DSBseq Quantitative Single-molecule sequencing of DSBs and repair products at three targets in tomato. A) Collection of time-course: mesophyll cell protoplasts are isolated from 2-3 week old seedlings of M82 *Solanum lycopersicum*. Duplicate samples are prepared with 200,000 protoplasts for each of the 7 time-points along 72 hours. CRISPR RNPs are introduced by PEG-mediated transformation. Samples are frozen at 0, 6, 12, 24, 36, 48, and 72 hours after RNP introduction and DNA is extracted. B) UMI-DSBseq target design: a primer specific to the target sequence, is coupled with a restriction enzyme site flanking the sgRNA target sequence in order to create an available end on intact molecules (WT or Indel) for ligation of adaptors. C) UMI-DSB library preparation: DNA extraction from time-course collection, containing WT (1), unrepaired DSBs (2) and intact molecules containing indels (3), is restricted *in vitro* with the restriction enzyme identified flanking the target cut site. Following end-repair by fill-in and A-addition, Y-shaped adaptors composed of P7 Illumina flow-cell sequences and containing i7 indexes and 9bp unique molecular identifiers (UMIs) are ligated to the unrepaired DSBs and restricted ends. Target-specific amplification by ligation-mediated PCR follows, with one primer identical to the adaptor sequence and containing the P7 Illumina tail (orange) and one primer specific to the target sequence (blue) with the P5 Illumina tail (red). This results in amplification of a single end of the DSB between the SpCas9 cut site and the primer. The red X represents the non-captured end of the DSB.

To address the challenge of directly quantifying the proportions of molecules, broken and repaired, we designed UMI-DSBseq, a ligation-mediated PCR-based assay combining a target-specific primer with a DSB flanking restriction enzyme site to capture both DSBs and intact molecules (Fig. 1B). The UMI-DSBseq assay facilitates high-resolution characterization of the state of all molecules by simultaneous ligation of adaptors containing unique molecular identifiers (UMIs) directly to unrepaired DSBs, as well as to intact molecules (both WT and containing indels) at the flanking restriction enzyme site, cleaved *in vitro* (Fig. 1B,C). To ensure capturing of all molecules in the pool including any potentially resected molecules, ends are repaired by fill-in of 3’ prior to adaptor ligation. The UMI-DSBseq protocol facilitates direct preparation of Illumina sequencing-ready libraries, enabling multiplexed sequencing of unrepaired DSBs in conjunction with repair products, all in a single tube (Fig. 1C). Following sequencing, molecules can be categorized as either unrepaired DSBs, WT intact molecules, or indel products of error-prone repair (www.github.com/daniebt/UMI-DSBseq).

### Capturing the kinetics of DSB induction and repair at endogenous loci

Three targets were selected for study in tomato, *Solanum Lycopersicum* cv. M82., at exons of three genes: *Carotenoid Isomerase, CRTISO* ^47^, *Phytoene Synthase 1, Psy1* ^27^ and *Phytochrome B2, PhyB2* (for oligos see Supplementary File 1). Timecourses consisting of 7 time-points, were collected for each target with 2 replicates independently transformed for each time-point (Fig. 2, for data see Supplementary File 2 \). Each sample collected for analysis contains a mixture of intact molecules (either uncut or repaired precisely), unrepaired DSBs and products of error-prone repair. Full negative-control time-course were collected for each target, consisting of 14 samples for each target without sgRNA, accounting for DSBs occurring due to DNA degradation or other artifacts (Fig. 2D-F, for data see Supplementary File 2).

**Figure 2.**
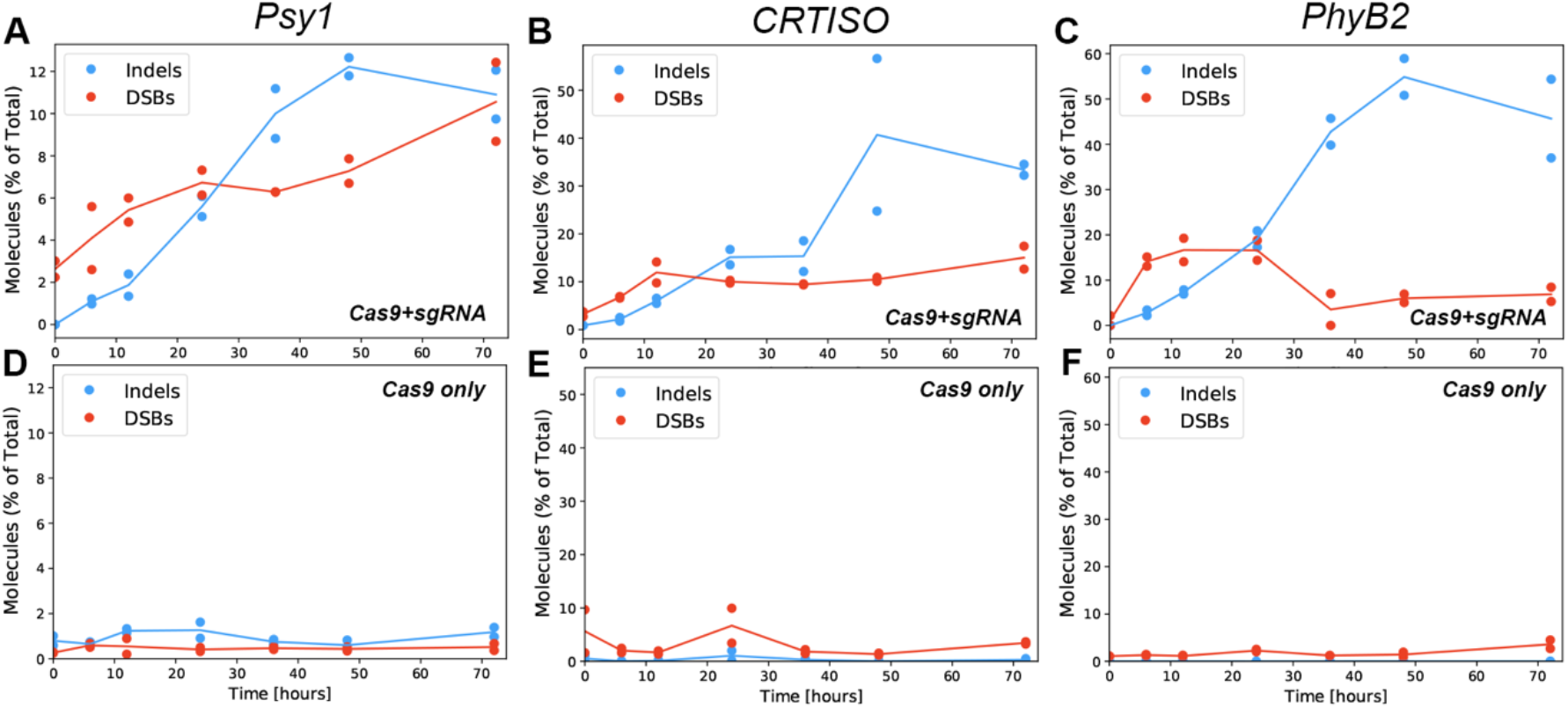
Patterns of Error-prone Repair along 72h time-course. A-F) Percent of molecules identified as unrepaired DSBs (red) and NHEJ-mediated indel (blue) out of total consensus sequences along 72h, (A-C) experimental and (D-F) control time-courses for *Psy1* (A,D), *CRTISO* (B,E), and *PhyB2* (C,F) with dots representing percent of molecules from each replicate and a line representing the mean of the duplicates. (see related: Figure S1, File S1)

Both unrepaired DSBs and indel-containing products of error-prone NHEJ can be detected at all three targets, with indel-accumulation peaking between 48 and 72 hours (Fig. 2A-C). DSBs first appear early in the time-course (Fig. 2A-C), with significant evidence at 6 hours in all 3 targets, compared to the controls (Fig. 2D-F). This suggests that RNPs are rapidly trafficked to the nucleus after transformation. Consistent with a longer repair process, molecules containing evidence of repair-error take hours to accumulate, with indels first detectable at low amounts at the 6-hour time-point and increasing following 24 and 36 hours (Fig. 2A-C). Out of the three targets analyzed, *PhyB2* displays both the highest frequency of indel products of repair-error and quantity of DSBs (Fig. 2C). While accumulating far fewer indels than *PhyB2, Psy1* has many detectable DSBs, reaching nearly 12% of the pool by 72 hours, approximately equal to the quantity of edited (indel) molecules (Fig. 2A). These results suggest that efficiency of SpCas9 DSB induction at this locus may be underestimated by examining editing alone.

Further insight into the dynamics of this process can be gained by evaluating the kinetics of the type of indels emerging from error-prone repair along the time-course. To this end, indels were categorized as either deletions, insertions, or deletions associated with 2 or more base pairs of flanking microhomology (Fig. S1). At the three targets, deletions rise rapidly (Fig. S1 A-F), particularly at *CRTISO* and *PhyB2* (Fig. S1 E-F). While, in contrast, insertions peak between 6-12 hours, before decreasing in total proportion, with this intriguing pattern most evident for *Psy1* (Fig. S1 D). Microhomology-associated deletions were far less abundant at the examined targets, emerging slightly later. While present in rather negligible amounts in *CRTISO* and *PhyB2*, these indels are more abundant in *Psy1*, likely due to more microhomologies flanking this locus (Fig. S1D-F, Supplementary File 1). A comparison of the results shown here (Fig. S1 G-L), to a previous analysis of indels obtained *in planta* with the same gRNAs at *Psy1* ^27^, or at *CRTISO* ^47^shows that most indels are similar. This suggests that the profile of indels is quite consistent, regardless of the cell types; and that the protoplast system is a good model for studying DSB repair in whole plants.

### Evidence of local off-targeting at *CRTISO*

At all three targets, the majority of DSBs are positioned around the expected SpCas9 cut site (between 3^rd^ and 4^th^ bps upstream of the PAM, between -3 and -4 positions, Fig. 3A-C). Alongside the presence of negligible DSBs in the negative control time-courses (Fig. 2D-F, Fig. 3A-C, Cas9 only) this evidence suggests that the UMI-DSBseq assay successfully captures Cas9-induced DSBs at the target sites. At *Psy1* and *PhyB2*, a sharp, single peak representing the vast majority of DSBs can be seen at the expected, SpCas9 -3 position, along with a small number of processed DSBs captured in close proximity to the target site (Fig. 3 A,C). In contrast, at *CRTISO*, two primary DSBs arise, with most captured ends positioned at the -4 position from the PAM (Fig. 3B).

**Figure 3.**
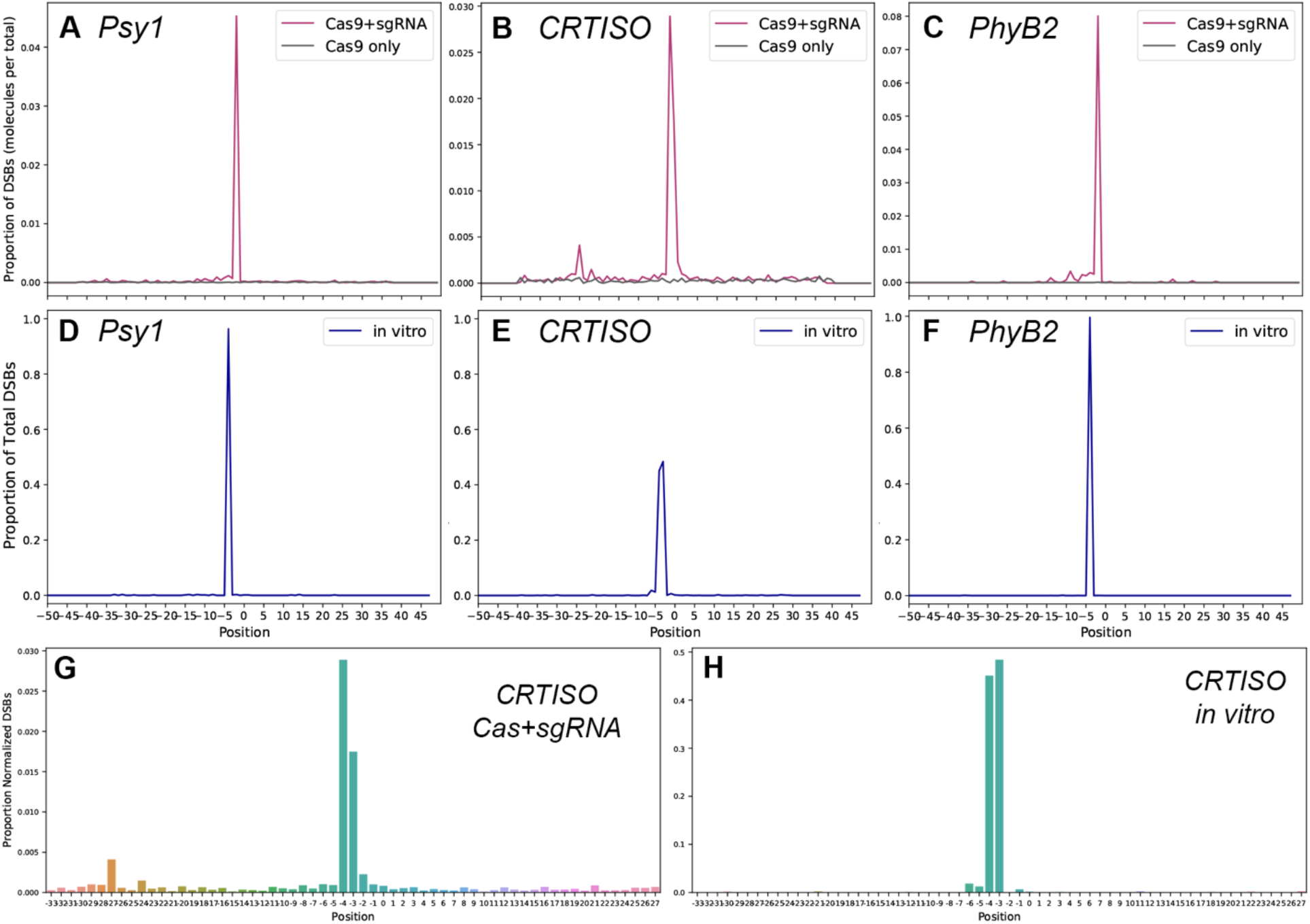
Local off-target at *CRTISO*. A-C) Proportion of total DSBs (normalized to total molecules) plotted by position along the target sequence with the expected cut site between -3 and -4 bp upstream of the PAM (A) *Psy1*, (B) *CRTISO* and (C) *PhyB2*. DSBs captured in experimental time-course in pink (Cas9+sgRNA, n=14) vs. control in black (Cas9 only, n=14). DE) Proportion of DSBs induced *in vitro* (normalized to total DSBs) plotted by position of the captured end at (D) *Psy1*, (E) *CRTISO* and (F) *PhyB2*, (n=4). G-H) Close-up of DSBs captured at CRTISO (B and E), plotted by position (G) time-course data vs. (H) *in vitro*.

In order to investigate whether these DSBs result from CRISPR-Cas9-mediated cleavage or from end-processing during repair, DSBs were induced in genomic DNA i*n vitro* and compared to those characterized in the experimental time-course (Fig. 3D-F, for data see Supplementary File 2). While at *Psy1* and *PhyB2 in vitro* induced DSBs occur precisely at the expected site, those captured at CRTISO, similarly to the timecourse results, appears to contain two peaks (Fig. 3E). A closer look confirms occurrence of the same two main peaks in the time-course as in the *in vitro* data, with differences in their relative abundance (Fig. 3 G,H). This supports that the two main -3 and -4 DBS termini are not due to *in vivo* end-processing but rather to local off-target cleavage by Cas9 at this locus.

### Kinetic model reveals efficient DSB induction coupled with precise repair

With the high-resolution data gathered, the rate of precise repair can be estimated using a set of ordinary differential equations, from the broken (DSB) and precisely repaired (intact) state, describing the curves of the different states over time, directly from the observed counts. We first implemented a simple 3-state model, with similar structure to that previously reported by Brinkmann et al ^45^, in which molecules are primarily classified as intact, DSB, and indels (Fig. 4). The rates of the flux between these different types of molecules are estimated as DSB induction (K_cut_), repair-error (E_repair_) and precise repair (P_repair_) rates, respectively representing the flux from intact to DSB, from DSB to indel, and from DSB back to intact (wild-type), in terms of proportion of molecules per hour (Fig. 4A).

Evaluating the flux through the intermediate, unrepaired DSB state, which is directly observed, facilitates decoupling the effect of induction efficiency from repair-fidelity at a given locus, compared to previous approaches ^45^. Using direct measurements of the state of each sequenced molecule in the pool (Fig. 2, Fig. 3, Supplementary File 2), we implement a maximum likelihood estimation of the model’s rates (Fig. 4, Table 1). This likelihood is maximized through extensive numerical optimization (Table 1); and uncertainty in the rate estimates is quantified through stratified bootstrap. Using this approach, the proportion of molecules flowing between the broken and repaired states can be estimated directly from the observations of single-molecule counts, revealing the rate of precise repair.

**Figure 4.**
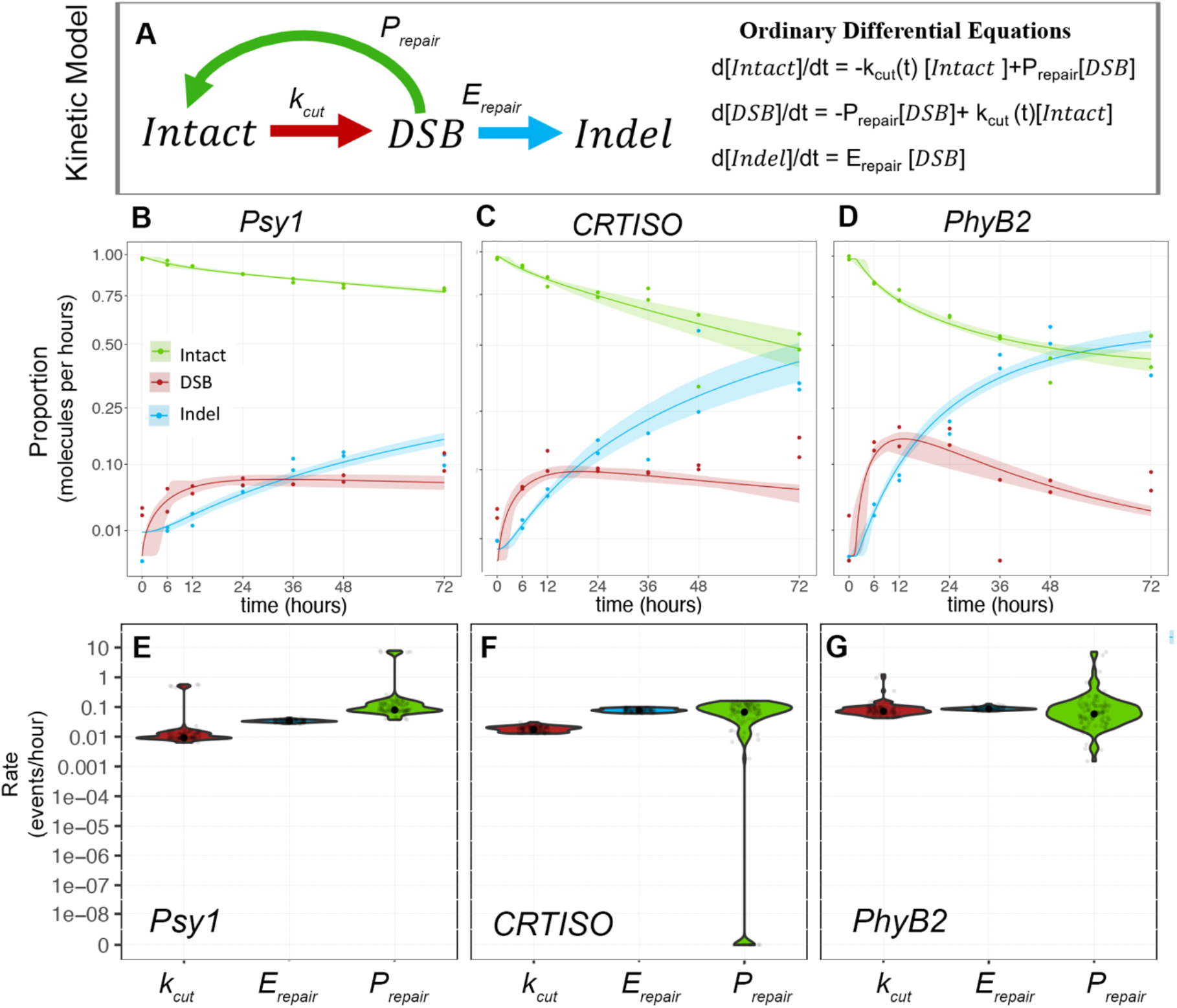
A 3-state model of DSB induction and repair. A) schematic and equations for a 3-state model of DSB induction K_cut_ in red, precise repair, P_repair_ in green and Error-prone repair E_repair_, in blue. B-D) Predicted fit of the model (lines) at *Psy1* (B), *CRTISO* (C), and *PhyB2* (D) for Intact molecules (green), DSB (red), and indels (blue). Confidence intervals are shown as shadings and calculated from 100 bootstraps of the data. Observed data are represented as dots. E-G) Rate constants estimated at *Psy1* (E), *CRTISO* (F), and *PhyB2* (G) in terms of number of events per hour per molecule. The smoothed distribution of the estimates obtained through the bootstrap procedure is shown as a violin plot. Mean estimate is shown as a black point. Grey points represent 100 instances of stratified bootstrap. (see related Table 1, Figure S2, Table S1)

**Table 1.** Rates and flow estimates for the 3-state model of DSB Repair.

| target | Process | rate constant<sup>a</sup><br>(proportion of total events/hour) | CI<sup>a,c</sup> [1%-99%] | flow<sup>b</sup> after 72h<br>(proportion of total molecules) | CI<sup>b,c</sup> [1%-99%] |
| --- | --- | --- | --- | --- | --- |
| *Psy1* | Cutting | 0.0092 | 0.0064-0.5434 | 0.5755 | 0.4031-31.0689 |
|  | Precise Repair | 0.0791 | 0.0373-7.2711 | 0.3581 | 0.1812-30.6583 |
|  | Repair Error | 0.0335 | 0.0276-0.0419 | 0.1518 | 0.1364-0.1683 |
| *CRTISO* | Cutting | 0.0175 | 0.0129-0.0290 | 0.8815 | 0.5794-1.3748 |
|  | Precise Repair | 0.0675 | 0.0000-0.1563 | 0.3799 | 0.0000-0.8654 |
|  | Repair Error | 0.0777 | 0.0611-0.0976 | 0.4375 | 0.3604-0.5178 |
| *PhyB2* | Cutting | 0.0718 | 0.0434-0.848 | 0.911 | 0.586 – 26.716 |
|  | Precise Repair | 0.0583 | 0.0036-4.747 | 0.3567 | 0.0229-26.0203 |
|  | Repair Error | 0.0864 | 0.0741-0.1220 | 0.5282 | 0.4966-0.5790 |

<sup>a</sup>Rates are reported as the number of events per molecule per hour. <sup>b</sup>The flow is reported as the proportion of molecules that experienced that event at the end of the experiment. <sup>c</sup>Confidence intervals (CI) are reported as the 1% and 99% percentiles of the estimates obtained from 100 stratified bootstraps of the data.

The fit of the model suggests that it offers a fair description of the process at the three targets along the time-course, with most observations falling within the range of values predicted by the model (Fig. 4 B-D, Table 1). Instead of imposing an induction curve measured *a priori*, by a proxy, our kinetic model directly estimates the induction curve from the data at each individual time-course, allowing for variations in genome search time and offering an approach more robust to mis-specification of the induction curve. The estimated induction curves are compatible between the three targets, suggesting relatively rapid induction within the early hours of the time-course with little decay within the 72-hour sampling window, consistent with the lack of depletion of DSBs in the observed time-courses (Fig. 2, Fig. S2, Table S1).

The estimated rate of DSB induction suggests that molecules are broken at a rate of approximately 1-8% per hour, with rate constant K_cut_ between 0.009-0.0718 of intact molecules per hour (Fig.4E-G, Table 1). These estimates indicate that in total, 57.5%, 88.2% and 91.1% of all molecules were successfully cleaved within 72 hours at *Psy1, CRTISO* and *PhyB2*, respectively (Fig.4E-G, Table 1). At the 3 targets, the rate of repair error, E_repair_, is estimated between 0.34-0.86 of DSBs per hour (0.0335, 0.0777, 0.0864 for *Psy1, CRTISO*, and *PhyB2*, Fig.4E-G, Table 1). These estimations predict that 15.2%, 43.8% and 52.8.% of the molecules present in the sample are repaired as indels over the time-course (Fig.4E-G, Table 1), consistent with the mean proportion of indels directly observed in the 48 and 72-hour sampling points at the three targets (Fig. 2A-C,Fig. S1 J-K).

Low indel accumulation coupled with relatively efficient cutting suggests the possibility that many of the broken molecules are repaired through high-fidelity mechanisms, leading to precise repair. Indeed, the rate of precise repair, P_repair_, is estimated between 0.058-0.079 molecules per hour at the three targets, suggesting that 35.81%, 37.99%, and 35.67% of all molecules have been broken and repaired precisely following the 72-hour time-course for *Psy1, CRTISO* and *PhyB2*, respectively (Fig.4E-G, Table 1). Statistical significance of precise repair is confirmed by 100 stratified bootstraps of the data for both *Psy1* and *PhyB2* (p-value < 0.01, Table 1). While for *CRTISO* the point estimate of the rate of precise repair (0.068) is comparable to those of *Psy1* and *PhyB2* (0.058-0.079), the same bootstrap approach could not rule out that precise repair is absent at this target, with confidence intervals 0-0.156 precise repair events per hour per molecule (Fig. 4E-G, Table 1). However, a model comparison of the 3-state model versus a model in which repair is only error-prone was significantly preferred for the full dataset at all targets (AIC-based relative likelihood < 10^-5^), as well as stably supported at *Psy1* and *PhyB2* in the bootstrapped data (100% of the bootstraps of *Psy1* and 96% of *PhyB2*, and 81% for *CRTISO*, Fig. S3, Table S2). Altogether, these results lead to the conclusion that DSB induction by the CRISPR-Cas9 RNPs is efficient; and is coupled with relatively high rates of precise repair.

### UMI-DSBseq captures the formation and processing of unrepaired DSBs

Captured DSBs may represent a pool of both newly formed breaks directly-induced by Cas9 (hereafter defined as directly-induced DSBs) as well as intermediates of active repair processes as they alter the DSB ends (hereafter defined as processed DSBs). Processed ends are the likely intermediates of indel formation upon repair, although in principle they can also be repaired precisely, e.g., through HR. In fact, the loss or gain of base-pairs at the ends resulting from errors in repair, would form transient intermediates offset from the expected position. Deletion intermediates would result in a shorter than expected captured DSB; while insertion intermediates (extended) or those of DNA synthesis (guide-side) would extend past the expected position (Fig. 5A). While DSBs captured at the targets are distributed around the expected break-site, processed DSBs were also detected at all targets while absent in the negative controls as well as in the *in vitro* samples (Fig. 3). To assess the dynamics of these putative intermediates, DSBs were categorized in directly-induced or processed DSBs depending on their position relative to the expected cut site (Fig. 5).

**Figure 5.**
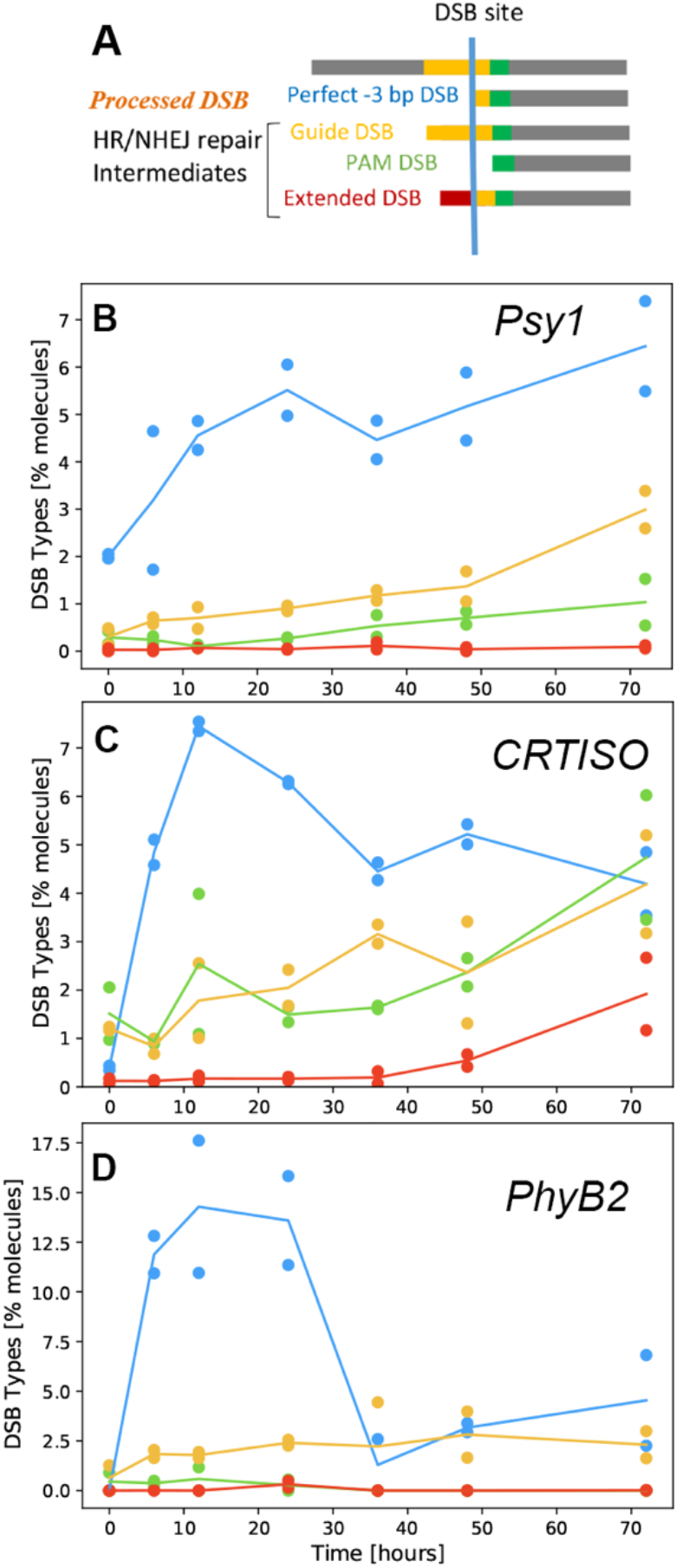
Characterizing putative intermediates of DSB repair. A) Schematic of categories of DSB types. Three categories of processed DSBs, Guide, PAM and Extended DSB. B-D) Percent of DSB types along the time-course of repair, for *Psy1* (B), *CRTISO* (C) (with perfect DSBs including both -3 and -4bp) and *PhyB2* (D). In blue perfect -3bp, yellow, guide-side DSBs, green, PAM-side DSBs and in red extended DSBs with extensions that do not match the target sequence.

Classifying DSBs in all three targets reveals that the majority of DSBs along the time-course are precise with no evidence of bp loss or gain (Fig. 5), suggesting they may be good substrates for the precise repair predicted by the 3-state model (Fig. 4, Table 1). However, at all 3 targets, some putatively processed DSBs can be identified, with classification revealing differing dynamics between the different types, compared to the perfect -3 DSB (Fig. 5B-D). The curve of precise -3 DSBs rises rapidly and begins depleting in both *PhyB2* and *CRTISO* (Fig. 5C,D). In contrast, at *Psy1*, the shapes of precise and processed DSBs are similar, with continued increase in abundance as the time-course progresses, perhaps suggesting evidence of continued cutting of precisely repaired molecules that have re-entered the pool (Fig. 5B).

At *CRTISO*, since two types of primary DSBs are observed and confirmed in the *in vitro* assay, in absence of the molecular machinery for DNA repair, we infer that both types of DSBs can be induced directly by Cas9, and so can be categorized as directly-induced DSBs (Fig. 3B,D,E,). Note, however, that the observed proportions of the two primary DSBs differ noticeably *in vivo* and *in vitro* (higher -4bp than -3bp *in vivo*, higher -3bp than -4bp *in vitro*) so that we cannot exclude that repair processes affect the two differently.

### Processed DSBs represent transient intermediates of error-prone repair

A dynamic model was built including ‘processed’ DSBs as a fourth state of the molecules, generated from directly-induced DSBs (Fig. 6, Table S3). In this model, the rates of the flux between these different types of molecules are estimated as DSB induction (K_cut_, from intact to direct DSB), processing of directly induced DSBs (K_processing_) and repair of both directly-induced DSBs (E_direct_, P_direct_) and processed DSBs (E_processed_, P_processed_) by error-prone and precise mechanisms. Using this approach, we can evaluate the contribution of DSB processing to high-fidelity and error-prone repair, estimating the flux through the intermediate state prior to re-ligation (Fig. 6). Altogether, the fit of the 4-state model makes predictions compatible with those estimated by the 3-state, with comparable values, and largely overlapping confidence intervals (Fig.4, Fig. S3, Table 1, Fig.6, Table S3). However, the 4-states model is significantly preferable (AIC-based relative likelihood < 10^-5^) over the 3-state model on the full dataset and in all 100 bootstraps of the data, at all 3 targets, suggesting it provides a better description of the complex dynamical process of DSB repair (Fig. S4G-I). Using this model, the rate of DSB processing and the type of repair at processed DSB intermediates can be estimated.

**Figure 6.**
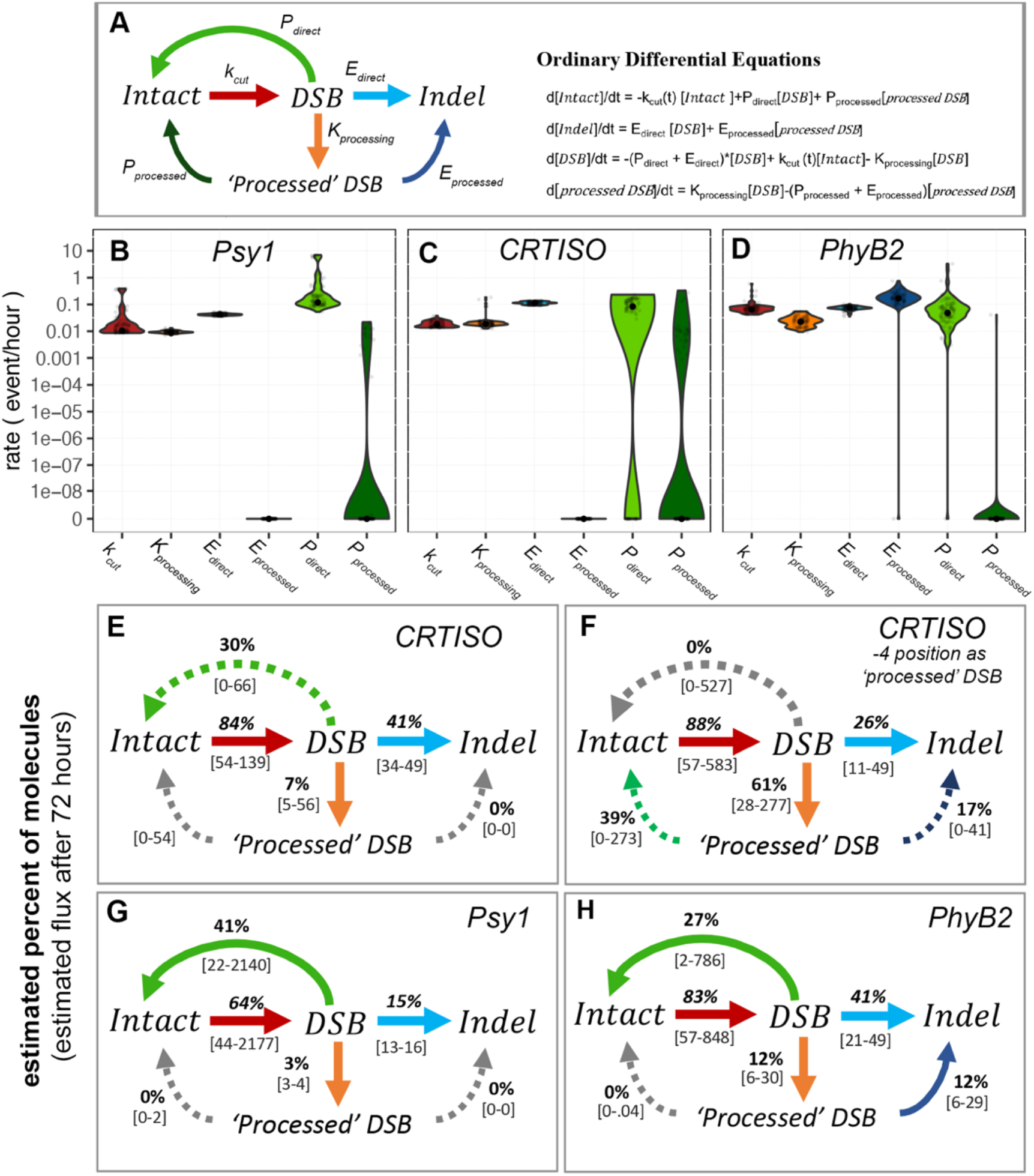
A 4-state model of DSB induction and repair, including processed ends. A) schematic and equations for a 4-state model of DSB induction, K_cut_, ends processing, K_processing_, repair from directly induced DSBs, namely precise repair, P_direct_ and error repair E_direct_, and from putative repair intermediates ‘processed’ DSB, including precise repair P_processed_ and error repair E _processed_. B-D) Violin plots of rate constants estimated at *Psy1* (B), *CRTISO* (C), and *PhyB2* (D) in terms of proportion per hour (as in Fig.4E-G). E-G) Schematic representation of the dynamic flow estimated in terms of percent of total molecules following 72 hours, at *CRTISO* (E) when the -4 position DSB is considered as directly induced (DSB) or (F) as ‘processed’ DSB, *Psy1* (G), and *PhyB2* (H). Grey arrows indicate rates estimated as 0, dashed arrows represent confidence intervals overlapping 0. CIs indicated in grey brackets and calculated from 100 iterations of boostrapping. (see related Figure S4, Table S2, Figure S5)

Consistent with their identification in the time-course data, processing of direct Cas9-induced DSBs is detected at all targets, accounting for 3-12% of total molecules along the time-course, with K_processing_ estimated between 0.009-0.0224 of directly-induced DSB molecules per hour (Fig.6B-D, Table S3). For *CRTISO*, two directly-induced DSBs (at positons -3 and -4) can be seen at *in vitro* and in the time-course data (Fig. 3 G and H) and it is possible that each could display different kinetics. This creates some ambiguity because the -4 bp cleavage can be considered either as directly-induced or processed end. Moreover, if one of the DSBs were staggered, as suggested in previous works^48,49^, such a break would produce a particularly good substrate for precise re-ligation.

In order to address this issue, we assessed *CRTISO* with the -4 local off-target DSB considered as ‘processed’ as opposed to directly induced (Fig. 6F,E), and compared the dynamics (Fig. 6E,F, Table S3, Fig. S5). This analysis reveals that most of the evidence for precise repair at this locus comes from DSB formed at directly at -4 (local off-target site) supporting the suggestion that indeed, it may represent a staggered cut formed by SpCas9 (Fig. 6E,F, Table S3, Fig. S5). Together, these results support a complex picture at this locus, with local off-target cleavage resulting in a two primary DSBs, with differing repair dynamics.

Precise repair of putative intermediates appears negligible at *Psy1* and *PhyB2*, with rate constant P_processed_ estimated at 0 (Fig. 6B,D, Table S3). At *Psy1*, only a small fraction of processed DSB can be detected in general; and they remain largely unrepaired (Fig. 6 B,G, Table S3). In contrast, processed intermediates at *PhyB2*, accounting for 12% of total molecules in the pool following 72 hours, are predicted to be repaired with error, at a rate of 0.1679 per hour (Fig. 6D,H, Table S3). These estimates suggest that all processed DSBs at this target represent transient intermediates of error-prone repair. Altogether, the high-resolution characterization of DSB intermediates facilitated by the UMI-DSBseq, coupled with a 4-state kinetic model further supports estimated rates of prominent precise repair while revealing additional hidden dynamics associated with repair-error, at *CRTISO* and *PhyB2*.

## Discussion

In this work we studied the kinetics of the DSB repair process using a novel molecular and computational toolkit, UMI-DSBseq. Starting with measurements of unrepaired DSBs and repair products, we characterized the dynamics of the repair process through kinetic modeling, successfully decoupling cutting efficiency from repair fidelity and revealing previously hidden aspects of both high-fidelity and error-prone repair (Fig. 7). These results establish that precise repair is a prominent feature of CRISPR-Cas9-induced DSB repair in somatic plant cells, and provide insight into the contribution of processed-ends to indel formation. Together, results shown here have numerous implications in both biotechnology and in the study of cellular processes involved in genome stability.

**Figure 7.**
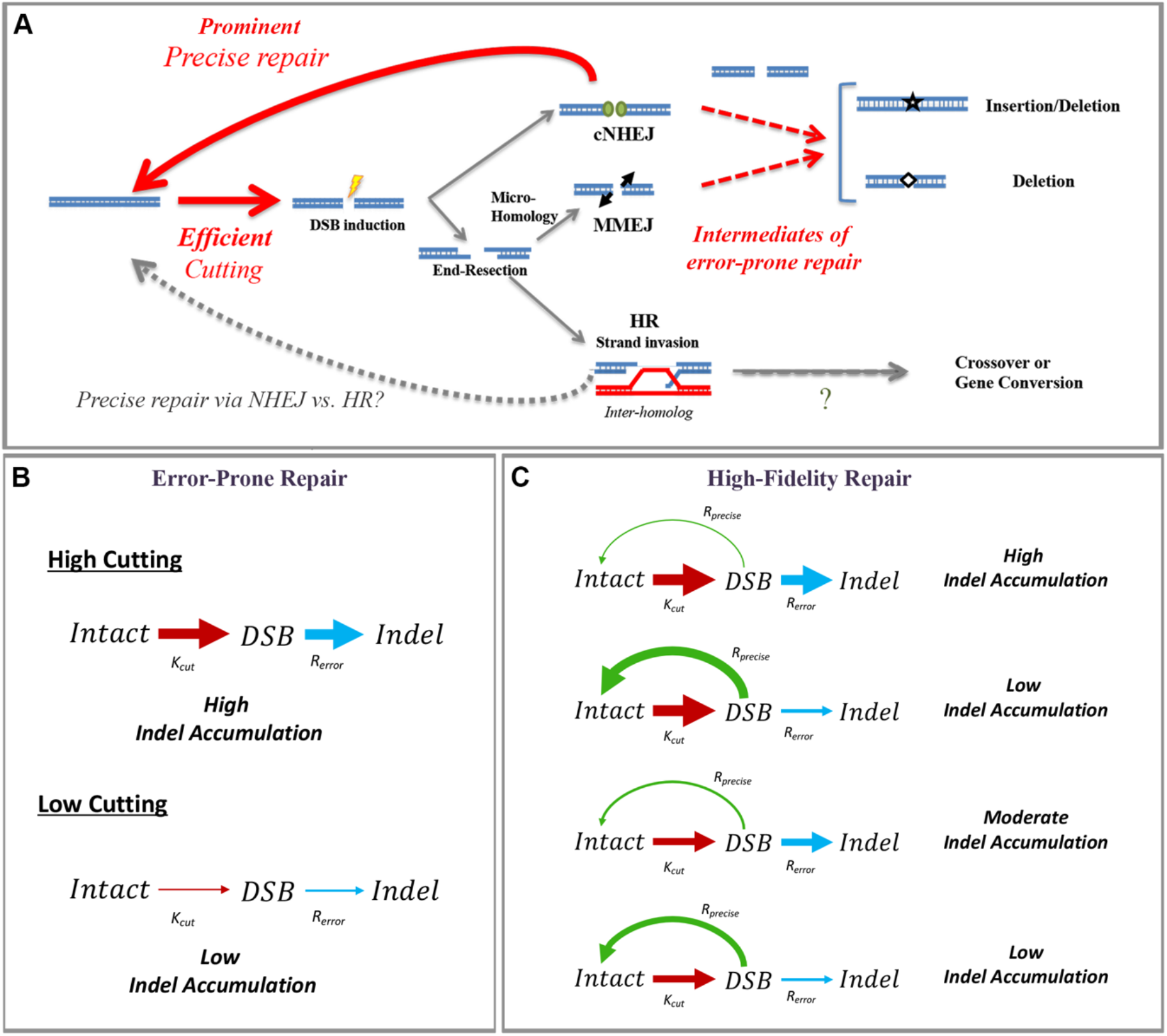
UMI-DSBseq coupled with likelihood-based modeling reveals previously hidden variables in the processes of DSB induction and repair. A) Decoupling of induction and repair revealing dynamics characterized by relatively efficient cutting and prominent precise repair. High-resolution characterization of unrepaired DSBs reveals intermediates of error-prone repair (*PhyB2* Fig. S1I). B) Model of DSB repair in the absence of precise repair was rejected by AIC relative likelihood. C) Models of DSB repair in presence of precise repair can explain the rate of indels formation at various loci (3-state is shown here for simplicity but same conclusion applies to the 4-state model). Thickness of the arrows indicates the efficiency, i.e. thicker arrow, higher proportion of molecules.

Deconvoluting DSB induction and repair-fidelity requires inferring the elusive flux from broken to intact molecules. This was achieved through precise measurements of intact, broken and repaired molecules, in a single assay. Compared to similar methods based on LM-PCR^45,50–56^, UMI-DSBseq facilitates simultaneous capturing of DSB intermediates alongside repair products, resulting in high-resolution quantification and characterization of all molecules along the time-course of repair. Moreover, the inclusion of UMIs enables a quantitative characterization of each repair intermediate and product through single-molecule sequencing. This has the advantage of allowing more direct measures of repair processes and to employ likelihood based statistical tools that can quantify and reduce our uncertainty. The UMI-DSBseq approach could be applied to a wide range of genome stability studies, such as the analysis of hotspots of recombination or transposition events, across a wide range of cell types.

At *PhyB2*, where evidence of processed ends is strongest, there is a nice correspondence between the position of primary indels and ‘processed’ DSBs characterized along the time-course (Fig.3C), suggesting that processed ends represent bona fide repair intermediates. Further insight into the high-resolution temporal dynamics at various loci, both from characterizing the unrepaired ends and probing repair kinetics, can help us better understand the factors impacting the type of pathway and errors that may emerge during repair. The 4-state model leads to the same conclusions as the 3-state model, namely, efficient DSB induction coupled with relatively high rates of precise repair. However, it adds a new dimension, facilitating exploration of complex dynamics, as in *CRTISO* (Fig. 6E-F, Fig. S5), and capturing the transition state following DSB formation, as with *PhyB2*, where processing of the ends contributes to error-prone repair (Fig. 5, Fig. 6H).

The new system presented here will be important for studying DSB repair. For example, the contribution of precise repair to the DSB repair process has been elusive so far. It might occur via precise NHEJ of unprocessed ends. In addition, precise repair could be achieved through HR, between sister chromatids or homologs. Recent works show evidence for the role of inter-homolog HR in DSB repair in somatic cells ^27,28,57^. Testing repair kinetics in NHEJ or HR mutants, or using DNA repair inhibitors, would enable to further dissect the mechanism of precise repair. Likewise, the effect of genes affecting end processing, either through deletion or addition of nucleotides, could be assessed using the 4-state model. These genes may contribute to error-prone repair or hinder the repair process leading to unrepaired ends and chromosomal rearrangements^58,59^.

Despite the wide-range use of CRISPR technology, the ability to faithfully predict efficiency of editing, or quantity of those predicted indels remains challenging ^41,60–62^. One likely reason for that, is that these studies have not decoupled the effect of cleavage from that of repair. In the absence of precise repair, the main factor limiting editing at a given locus would be the cleavage efficiency of the targeted endonuclease, namely the sgRNA and Cas9 protein (Fig. 7B). However, we found that cleavage efficiency was high at all targets, including in the low-editing Psy1 locus, showing that ignoring precise repair cannot explain the experimental data, and introducing precise repair in the 3- and 4-state models provided a much better description of the process. Therefore, our results suggest that CRISPR editing efficiency may vary, due to the combined effect of precise repair, ends processing and error-prone repair (Fig. 7C) and to a lesser extent due to cleavage efficiency. This complexity, involving multiple components of genome maintenance machineries, explains why it has been difficult to predict CRISPR efficiency.

Our work decouples the various processes involved in DSB formation and repair, opens new possibility to bridge the gap between the basic science of DNA repair and the application of the CRISPR technology. Altogether, this reinforces the need for further study. For example, large screens are identifying mutations in DNA repair proteins that increase efficiency of editing across multiple platforms ^63,64^. Further assignment of specific genes to the different stages in the DSB repair process (induction, processing and fidelity of repair) can provide additional insight into optimizing this technology across a wide range of organisms. In general, by facilitating multiplexed single-molecule sequencing of repair intermediates and products, we open the door for application of likelihood-based statistical methods to evaluate the process DSB induction and repair across diverse species, cell-types, and genomic contexts.

## Acknowledgements

We wish to thanks the Israel Science Foundation ISF2332/19, and the ministry of innovation CRISPR-IL consortium to AAL for financial support and the Clore Scholars Fund for the PhD fellowship for DBT; Dr. Barry Cohen for his assistance in optimizing the protoplast protocol; the protein expression unit at the Weizmann Institute for the Cas9 protein purification. We would like to thank also Maayan Guetta, Ilan Hadad and Ido Sela for their technical assistance.

## Author Contributions

This work was supervised by AL and CMB. DBT and CMB conceptualized and designed the UMI-DSBseq molecular assay. UMI-DSBseq analysis pipeline was written by DBT and AC. DBT, AH and CMB calibrated and optimized the RNP transformation protocol. DBT conducted experiments, library preparation and UMI-DSBseq analysis. Sequencing runs were carried out and managed by CMB. All kinetic modeling design, optimization, and implementation along with all statistical analysis was carried out by FM. Manuscript was written by DBT, FM, and AL.

## STAR Methods

### RESOURCE AVAILABILITY

#### Lead Contact

Further information and requests for resources and reagents should be directed to and will be fulfilled by the lead contact

#### Data and Code Availability

BASH scripts for calling consensus sequences and Jupyter notebooks for characterizing DSB and repair types for UMI-DSBseq data can be found at https://github.com/daniebt/UMI-DSBseq. Kinetic model can be found at www.github.com/fabrimafe/DSBtimecourse

### EXPERIMENTAL MODEL AND SUBJECT DETAILS

#### Plant Materials and Growth

M82 Tomato seeds were gas sterilized and sown on Nitsch media in magentas under long-day conditions, at 23 C.

### METHOD DETAILS

#### Protoplast Isolation and Purification

The protocol was adapted from Yoo et al. ^65^. Briefly, first true leaves of 14 to 20-day-old seedlings are sliced into thin strips and immersed in 15ml of enzyme solution (3.65g Mannitol, 2ml 0.5M MES pH=5.7, 500 ul 2M KCl, 0.75g Cellulase R10 Duchefa, 0.2g Macerozyme Duchefa, 500ul 1M CaCl2, 0.05g BSA) in a Petri dish, and incubated in the dark overnight with mild shaking (25 RPM) for 14-16 hours. Extracted protoplasts strained through a 0.1 um filter and washed with W5 solution (154 mM NaCl, 125 mM CaCl2, 5 mM KCl, 2 mM MES pH=5.7). Healthy, intact protoplasts are isolated using sucrose gradient (23%, 11.5 g per 50ml). Following additional washing with W5, protoplasts are diluted to a concentration of 1 million cells/ml MMG solution (0.4M Mannitol, 15 mM MgCl2, 4 mM MES pH=5.7).

#### sgRNA preparation

sgRNAs are purchased from IDT as tracrRNA and target-specific crRNA. Both are diluted in TE buffer to a concentration of 200uM each. Equimolar ratio of the two components are mixed with Nuclease Free Duplexing Buffer and heated at 95C for 5 minutes before slowly cooling to room temperature. Stored at -20 C until day of transformation.

#### SpCas9 protein expression

SpCas9 protein was expressed from pET-28b-Cas9-His^66^ (Addgene plasmid # 47327 RRID:Addgene_47327) by the proteomics Unit at the Weizmann Institute.

#### Preassembly of CRISPR RNP

For each sample 10ug of SpCas9 and 20ug of duplexed sgRNA are mixed with NEB Buffer 3.1 at 20 ul total volume and incubated at room temperature for 15 min. RNPs are prepared immediately prior to transformation.

#### PEG-mediated Transformation of CRISPR RNP

20 ul preassembled RNP is added to 200,000 protoplasts in 200 ul MMG solution. An equal volume of PEG solution (40% [w/v] PEG 4000 Sigma, 0.2M Mannitol, 0.1M CaCl2, DEPC treated water) is added and mixed gently prior to 20 min incubation in the dark. Then the samples are diluted with W5 solution and incubated for 15 min in the dark. The samples are centrifuged at 450xg with soft start/stop and resuspended in 1ml WI solution (0.5M Mannitol, 20 mM KCl, 4 mM MES pH=5.7) before incubation in the dark. At each time-point, duplicate samples are independently transformed, for 14 samples per time-course, for hours 0, 6, 12, 24, 36, 48, 72 hours post transformation. The samples for time 0 are frozen prior to addition of the WI, with the time of freezing set to the start of the time-course.

#### Sample collection and DNA extraction

Samples are pelleted using centrifugation at 450xg, WI is removed and samples are flash frozen in liquid nitrogen. DNA is extracted using NucleoSpin Plant II extraction kit (Machery-Nagel Cat #) with modified protocol. Briefly, 600 ul PL1 lysis buffer are added to each sample with 10 ul RNAse and incubated 1 hour at 65°C. After running through column, 675 ul PC buffer is added. Washing is done with 2x with PW2 buffer before elution in 50 ul elution buffer.

#### UMI-DSBseq Adaptor Design and Preparation

Design of the UMI-DSBseq adaptors was adapted from on the P7 tail of the xGen UDI-UMI adaptors from IDT (https://eu.idtdna.com/pages/products/next-generation-sequencing/adapters/xgen-udi-umi-adapters), containing an 8bp index for barcoding from the IDT8_i7 index list, and a 9bp unique molecular identifier, for analysis with the recommended pipeline for single molecule sequencing. Two oligos are ordered from IDT as ULTRAMERs, a long tail containing P7 tail with UMI indexes, along with a final T at the 3’ end with a phosphorothioate bond (*T) for sticky T/A ligation, as well as a semi-complementary oligo (P7 forward tail: GATCGGAAGAGCGGGGACTATTTGC). The two oligos are annealed by heating to 95°C and allowing to cool to room temperature. Prepared adaptors are diluted to 1uM in 10mM Tris pH=8, and stored at -20°C until use.

#### UMI-DSB Library Preparation

20 ul of extracted DNA (25-50 ng) is restricted overnight with target-specific enzyme at 37°C. After cleaning with magnetic beads, restricted DNA is end-repaired using DNA Polymerase 4. A-addition using Klenow is followed by UMI-DSBseq adaptor ligation using DNA Ligase 4 with 2 ul from each adaptor, in Quick Ligase buffer with cleaning between each step. Target-specific amplification is achieved using 1 primer specific to the target sequence (see table below) with a tail composed of the P5 illumina tail, and 1 primer identical to the 5’ end of the P7 adaptor tail sequence from the UMI-DSBseq adaptors (CAAGCAGAAGACGGCATACGAGAT)^67^. The adaptor specific primer can bind only loci for which second strand synthesis was already achieved due to the elongation from the target-specific primer. In the final step, i5 indexes are added using an additional PCR with the short enrichment primers used in TruSeq Illumina library preparation. Following cleaning, concentration is measured using Qubit (Cat #) and evaluated for quality and size using TapeStation

#### Next-Generation Sequencing of UMI-DSB libraries

UMI-DSBseq libraries are sequenced using 150bp paired-end Illumina Nextseq or Novaseq kits. Libraries were sequenced with either NextSeq or Novaseq Illumina machines at the Weizmann Institute using 300 bp paired-end kits. The run settings include 17 bp index 1 and 8 bp index 2, with 149-151 bp for each read 1 and read 2. Libraries are individual time-course samples are sequenced with 1-5 million reads per library.

#### Control Time-courses

For each target, full control time-courses were developed either by transforming Cas9 only to WT protoplasts or by mock-transformation Cas9 expressing protoplasts. Processing proceeds as described above.

#### In-vitro DSB assay

Genomic DNA extracted from tomato protoplasts, equal to the quantity processed for samples in the experimental time-courses (40% of sample DNA from 200,000 protoplasts), is cleaved in-vitro with 40% of the RNP produced as described above – Following incubation at 37°C for 1 hour, EDTA is added to stop the reaction, and the cleaved gDNA is cleaned using magnetic beads, followed by processing through the UMI-DSBseq workflow, as described above.

### QUANTIFICATION AND STATISTICAL ANALYSIS

#### Demultiplexing and generation of consensus sequences

Demultiplexing is done using bcl2fasq, splitting the index 1 read into the UMI file and index file. Data processing pipeline was adapted from the IDT pipeline (https://www.youtube.com/watch?v=68sca_jsqg8&ab_channel=IntegratedDNATechnologies) for building consensus sequences using FGBIO (https://github.com/fulcrumgenomics/fgbio).

Fastqs are aligned to a reference of the target sequence using bwa-mem (bwa mem -t 8 $reference $fastq_R1 $fastq_R2 > $mapped_bam). Unmapped BAM files are generated using picard FastqToSam. The unmapped and mapped BAM files are merged using picard MergeBamAlignment (MAX_GAPS=-1, CLIP_ADAPTORS=true) and annotated with UMIs using FGBIO. Reads are grouped by UMI using FGBIOs GroupReadsByUmi (strategy=adjacency, edits=1, min-map-q=0,assign-tag=MI). Finally, consensus sequences are generated using FGBIO CallMolecularConsensusReads with min-reads=2 and minimum input base quality set to 20. Final BAM files with consensus reads are converted to Fastqs using SamToFastq from picard, and joined using ea-utils fastq-join.

#### Quantification and Characterization of Consensus Sequences

The joined fastq files are processed through the UMI-DSBseq analysis pipeline (https://github.com/daniebt/UMI-DSBseq), and available as Jupyter notebook. The workflow requires as input the joined fastq files of the consensus sequences, and a sample sheet in .xlxs format containing the file name, sgRNA sequence, primer sequence, and amplicon sequence as well as relevant run information, for each sample in the time-course. *The workflow proceeds as described in the following sections*

##### Extracting WT Reference sequence from sample sheet

A 100 bp reference sequence (WT_REF) is extracted from the sample sheet, 50 bp upsteam or downstream of the DSB site for each target. Fastq files are uploaded using BioSeq SeqIO package, outputting dictionary with table of reads for each sample.

##### Determining the State of the read as Intact, Indel containing, or DSB

12bp indicators on each end of the 100 bp reference window are aligned to each read, using pairwise2 local alignment from the BioSeq package. A molecule is defined as Intact, if both left indicator and right indicator receive an alignment score of at least 10/12 bp. For each Intact molecule, the sequence between the two indicators (inclusive) is extracted for further analysis (seq_window). A molecule is defined as a DSBs, if only the left indicator receives the minimum alignment score. In the case of defined DSBs, region is extraction from the left indicator (inclusive) to the end of the reads (seq_window). Any reads that do not either criteria for Intact or DSB, or that contain more than 4 ‘N’s in their sequence are defined as NA. PCR contamination is filtered by by removing any molecules ending in a known primer sequence as opposed to the restricted end.

##### Characterizing footprints of repair error

seq_window for each read is aligned to the 100 bp WT_REF using pairwise local alignment from the BioSeq package. Intact reads are defined as WT if no gap was opened in either the read or the reference. If a gap is opened in the WT_REF, the type of the molecule is defined as Insertion, and the state is changed to NHEJ. If a gap is open in the seq_window of the read, the Type of the molecule is defined as deletion and the state is changed to NHEJ. The name of the indel is defined by the missing or added bases prefixed by ‘+’ or ‘-‘. Deletions associated with microhomology at the site, are defined as MH_Del, and the identity of the microhomology is reported, defined as 2 or more bp of microhomology flanking the deletion.

##### Characterizing DSBs by type and position

the seq_window is aligned as with the Intact molecules, and ‘N’s at the end of the sequence are trimmed. Based on the size of the DSB, they are defined as precise, at the position 50 in the WT_REF, or either guide-side or PAM-side, if they are positioned to either side. In addition, if the DSB is longer than expected and contains non-matching sequences to the reference, it is categorized as ‘Extended’. Characterized reads are grouped and counted.

#### Kinetic Model

A set of ordinary differential equations adapted from Brinkman et al.,^45^ were used to develop a model for calculating the rate constants between the different states in the process. The model is fit to the data using gradient descent optimization of the loglikelihood using the R package opmitx. To constrain the optimization procedure we imposed a maximum cutting rate of 10 cuts per hour, which largely exceeds the highest estimates for CRISPR-Cas9. To ensure that we converge to the global maximum, we initialize the optimization from 5.000 points sampled randomly following an exponential distribution with average 0.01 and within the constrained range of potential values. The log-likelihood is calculated assuming independence between molecules within each sample, so that the proportion of sampled molecules of the different types is expected to be proportional to the total proportion of molecules present and expected from the dynamical model. This results in a multinomial sampling, for which the likelihood can be computed exactly.

To control for errors resulting from sequencing or other steps of the UMI-DSBseq assay we used the control time courses (in which only intact molecules are expected) to estimate the probabilities that an intact molecule is classified as belonging to a different state. We then built error matrices specifying the probability that a molecule of any state is classified as that of another state under the assumption that intact molecules carrying indels would show the same error rates as intact molecules carrying the original sequence, and that the chance of a broken molecule to be read as intact is negligible. Note that the proportion of broken molecules is usually much lower than that of intact ones, so that the latter assumption is expected to have only minor effects. We explored other assumptions observing negligible effects. Such different implementations of the error matrix can be explored using the -e flag in the provided code. Additionally, for the 3-state model imprecise DSBs are modeled as errors arising from precise DSBs. Since precise DSBs are not found in the control time courses, the probability that such errors occur is estimated dynamically as an additional parameter in the optimization of the likelihood of each time course. The code for the model estimation can be found at https://github.com/fabrimafe/DSBtimecourse.

#### Determining the parameters of the induction curve

The accessibility of Cas9 and proportion of active RNPs are modeled by assuming that the rate of cutting depends on K(t), which is a function of time t. We chose the function:

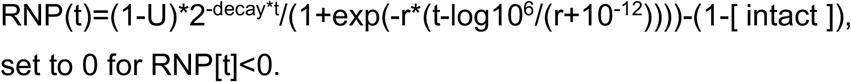

This function describes a constant decay of availability of RNPs, with half-life 1/r2, a logistic induction with slope r0 and constrained to assume value 10^-6^ at time 0, and implies that only intact molecules can be cut.

#### Calculating confidence intervals

Confidence intervals are calculated using a stratified bootstrap approach: 100 simulated datasets were generated by resampling the original data-points with replacement within each time point; for each of these datasets the optimization procedure was repeated and the 1% and 99% percentiles of each parameter estimates used as confidence intervals.

#### Model comparison

The Akaike information criterion was used for model comparisons. The relative likelihood was calculated from the AIC. To assess the stability of the model comparison, AIC and relative likelihood were computed for all bootstrap iterations. A proportion higher than 95% of the bootstraps supporting the more complex model, i.e. with relative likelihood lower than 1 and lower AIC value for the more complex model, was used as a significance threshold.

#### 3-state model

We implemented and compared two different dynamical models with different degree of complexity and detail, differing in the rates being modeled and in how DSBs ending at positions corresponding to the expected Cas9 cutting site and other unexpected DSBs, defined as precise and imprecise DSBs, respectively, are treated.

The *3-state model* follows the dynamics of intact molecules, DSBs and indels, without considering any biological difference between DSBs ending at positions corresponding to the expected Cas9 cutting site (precise DSBs) and other DSBs (processed DSBs). Specifically, unexpected DSBs are modeled as errors introduced by the experimental processing or sequencing which result in some DSBs to end at positions different than that corresponding to the expected Cas9 cutting site. Hence, precise DSBs are observed as imprecise DSB with a given probability p_error_. The systems of ordinary differential equations describing these models are shown below. The 3-state dynamical model can be represented as:

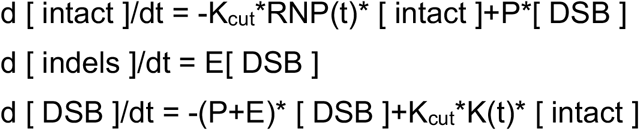

where K_cut_ is a constant cutting rate, P and E are the repair rates of DSBs towards intact molecules and indels, respectively. At each time t, the proportion of processed DSB is simply estimated as a proportion perror of the DSB. Hence:

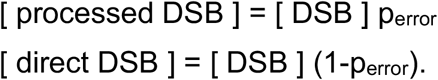

For the 3-state model without repair we used the same model by fixing P to 0.

#### 4-state model

A *4-state model*, which distinguishes between precise and imprecise DSBs. This more complex model, allows imprecise and precise DSBs to have different dynamics. Imprecise DSBs can appear through the successive molecular processing of precise DSBs. Both direct and processed DSBs can undergo precise repair to intact molecules (P) or error-prone I to indels, with rates P_direct_ and E_direct_, and P_processed_ and E_processed_, respectively. Direct DSB are converted to processed DSB with rate K_processing_.

The 4-state model is:

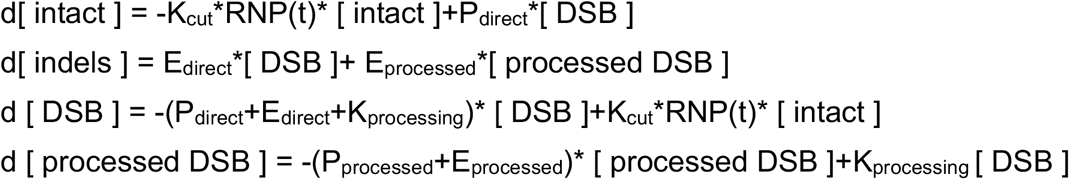

We also analyzed variations of this model, allowing for imprecise DSBs to arise because of off-target cutting events by Cas9, with a two cutting rates, K_direct_ (referred to as Kcut for other models) and K_processed_ (SIXX).

#### Calculating Flow over 72 hours

For each rate we calculated the amount of molecules out of the ones initially present in the pool which underwent a specific transformation, e.g. from intact to precise DSBs with rate K_cut_, through numerical integration of the systems of ordinary differential equations. Note that the total amount of molecules going through each rate can exceed 100%, e.g. more intact molecules than present at the beginning of the experiment can be cut to precise DSBs over the rate of the time course because some intact molecules are replenished via perfect repair. Numerical integration was computed at a resolution of 500 intervals each hour of the time course. The code for the numerical integration is deposited as an R script at github.com/fabrimafe/DSBtimecourse/plot_bootstraps.R

## KEY RESOURCES TABLE

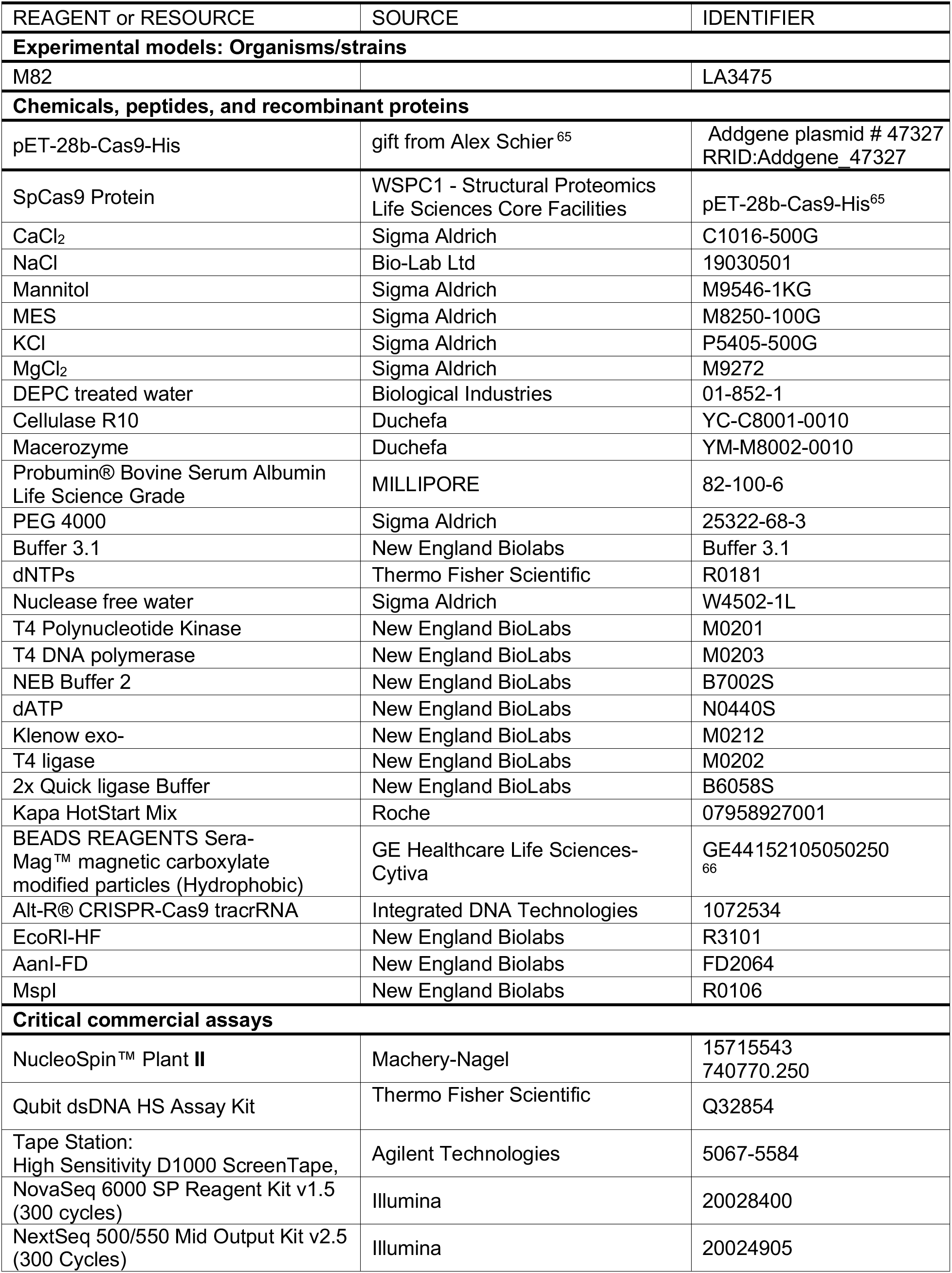
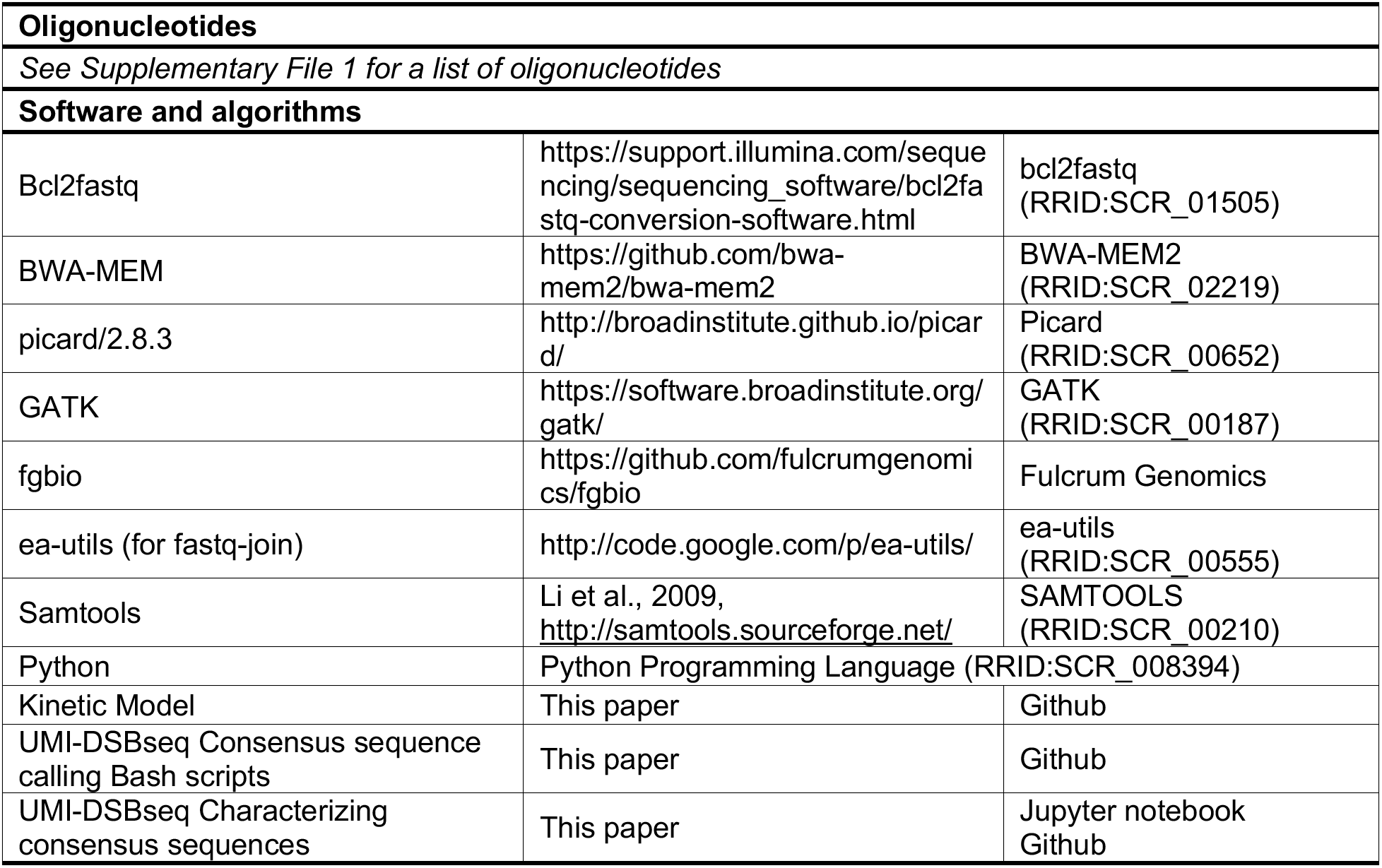

## Supplementary Figures and Tables

**Figure S1.**
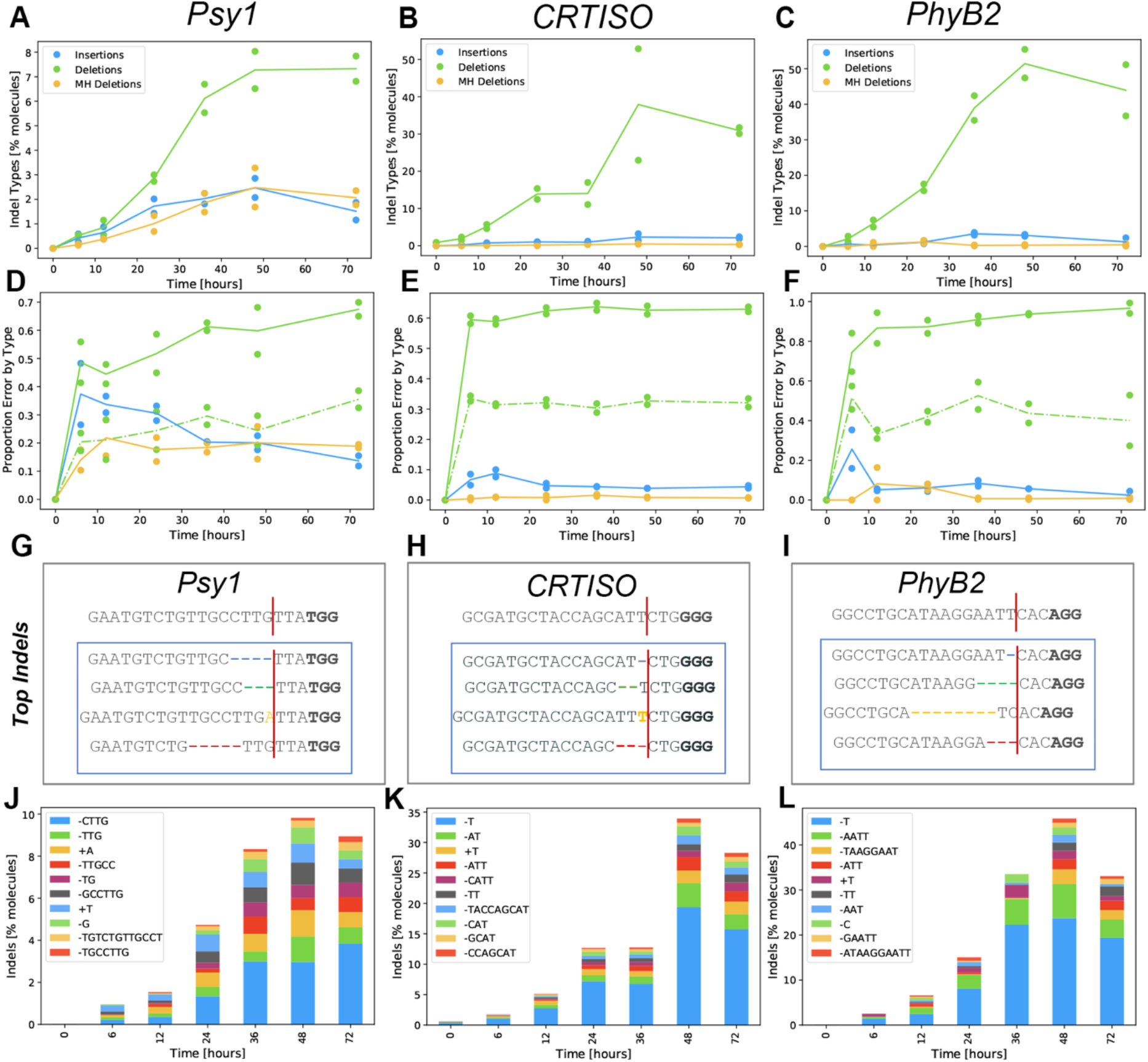
Footprints of error prone repair at 3 targets in tomato. A-C) Percent of indel containing out of total molecules for (A) *Psy1*, (B) *CRTISO* and (C) *PhyB2*. D-F) Proportion of each type out of total indels, during the time course from 0 to 72 hours, for (D) *Psy1*, I *CRTISO* and (F) *PhyB2*, either insertion (blue), deletions (green) or deletions associated with 2 or more bp of microhomology (MH Deletions, yellow). Dots represent each duplicate and lines represent the mean of the replicates. G-I) Sequence of each target with PAM indicated in bold and red line for the expected cut site. Top indels represented in the box, ranked in order of abundance for (G) *Psy1*, (H) *CRTISO*, and (I) *PhyB2* J-L) Top 10 indels at each timepoint, stacked bar plots of the mean of the two replicates. (J) *Psy1*, (K) *CRTISO*, and (L) *PhyB2*.

**Table S1.** Estimates of the Error and Induction parameters for the 3-state model of DSB Repair.

| target | Parameter | rate constant <sup>a</sup><br>(proportion<br>events/hour) | CI <sup>b</sup> [1%-99%] |
| --- | --- | --- | --- |
| *Psy1* | pError | 0.2052 | 0.1659-0.2392 |
|  | r [x10<sup>-3</sup>] | 9.4021 | 0.0025-20.9090 |
|  | decay | 0.0000 | 0.0000-0.0000 |
|  | U | 0.0000 | 0.0000-0.0001 |
| *PhyB2* | pError | 0.0817 | 0.0480-0.1353 |
|  | r [x10<sup>-3</sup>] | 0.0080 | 0.0029-1.0793 |
|  | decay | 0.0000 | 0.0000-0.0073 |
|  | U | 0.4047 | 0.0000-0.4395 |
| *CRTISO* | pError | 0.3200 | 0.2332-0.3908 |
|  | r [x10<sup>-3</sup>] | 0.0488 | 0.0053-24.2933 |
|  | decay | 0.0000 | 0.0000-0.0048 |
|  | U | 0.0000 | 0.0000-0.1179 |

<sup>a</sup> Rates are reported as the number of events per molecule per hour, with the exception of the rate of the steepness of the induction curve (r induction), which is reported as the number of events in 0.001 molecules. <sup>b</sup> Confidence intervals are reported as the 1% and 99% percentiles of the estimates obtained from 100 stratified bootstraps of the data.

**Figure S2.**
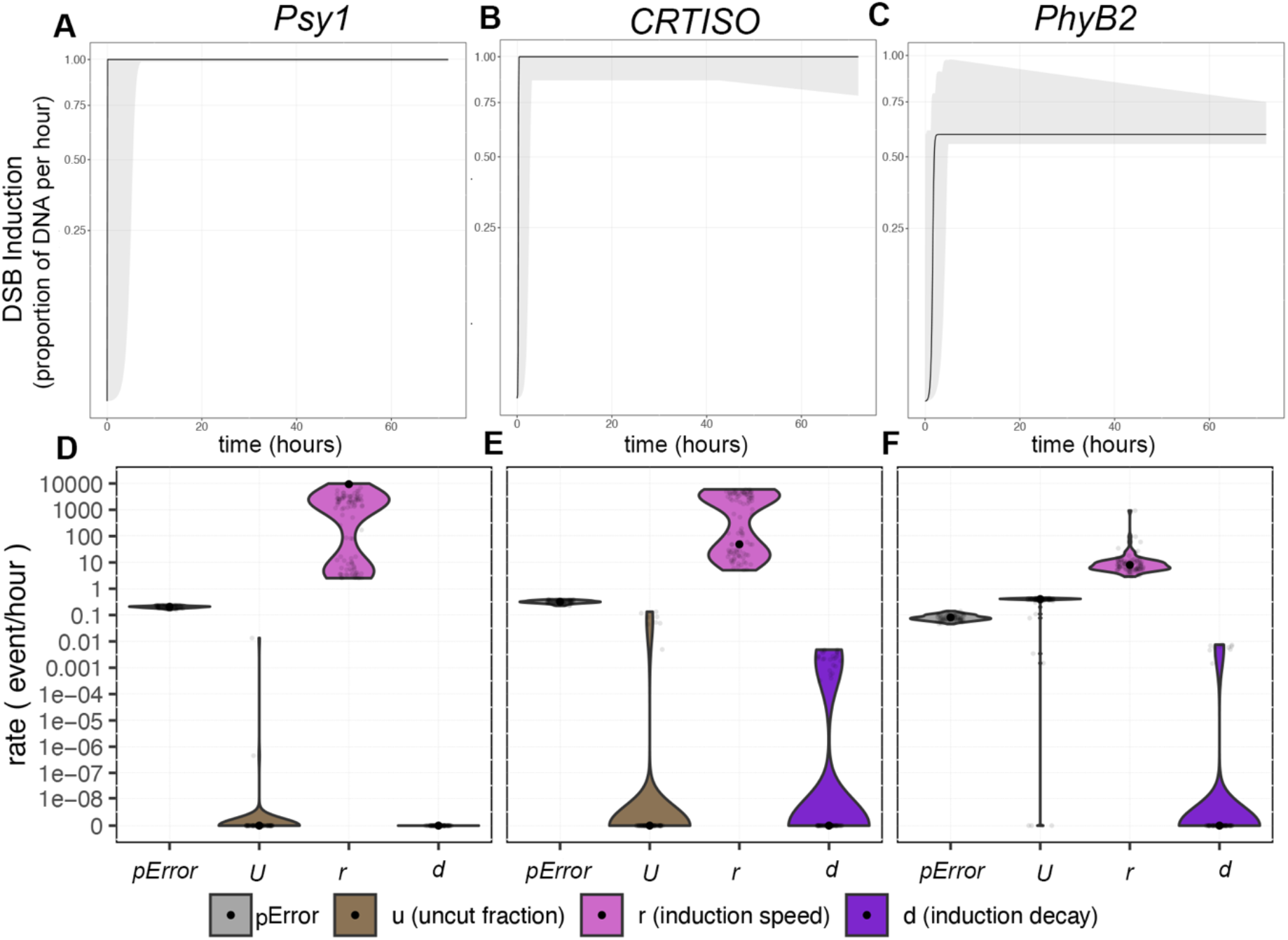
Induction Curve predicted by 3-state model. A-C) Induction curves with confidence intervals from the bootstrap indicated in gray shading D-F) Estimates for Error (unexpected DSBs modeled as errors introduced by the experimental processing or sequencing which result in some DSBs to end at positions different than that corresponding to the expected Cas9 cutting site) and the three parameters of the induction curve, uncut fraction (u), induction speed (r), and induction decay (see Methods). The uncut fraction (u) represents the estimated proportion of intact DNA molecules which are not being cut throughout the time-course. The induction speed (r) determines the slope of the induction curve, i.e. how rapid is the increase in cutting rate after the delivery of the RNPs, while the induction decay determines the rate at which the induction of DSBs decrease over time, possibly as a result of depletion of RNPs. The smoothed distribution of the estimates obtained through the bootstrap procedure is shown as a violin plot. Mean estimate of 100 iterations of bootstrapping is shown as a black point at A,D) *Psy1*, B,E) *CRTISO* and C,F) *PhyB2*. (D,F). (see related Figure 4, Table1, Table S1)

**Figure S3.**
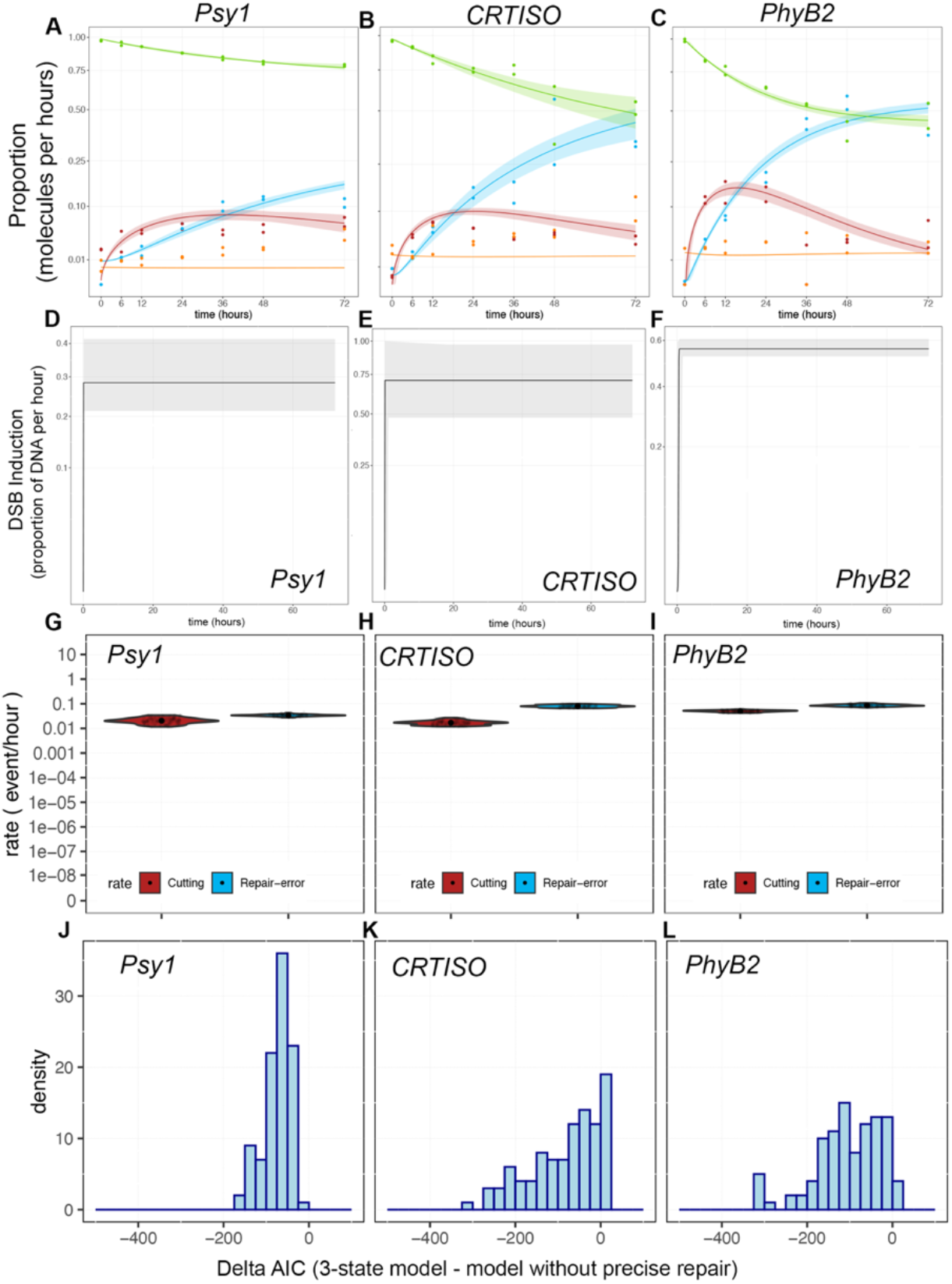
Kinetic model with no precise repair. A-C) Predicted fit of the model (lines) at *Psy1* (A), *CRTISO* (B), and *PhyB2* (C) for Intact molecules (green), DSB (red), indels (blue) and DSB Errors (orange). Confidence intervals are shown as shadings and calculated from 100 bootstraps of the data. Observed experimental data represented as dots. D-F) Induction curves for *Psy1* (D), *CRTISO* (E), and *PhyB2* (F) with confidence intervals from the bootstrap indicated in gray shading. G-I) Rate constants estimated at *Psy1* (G), *CRTISO* (H), and *PhyB2* (I) in terms of number of events per hour per molecule. The smoothed distribution of the estimates obtained through the bootstrap procedure is shown as a violin plot. Mean estimate is shown as a black point. Grey points represent 100 instances of stratified bootstrap. J-L) Difference in AIC (Delta AIC, where AIC stands for Akaike Information Criterion) between the 3-state model with and without perfect repair for Psy1 (J), *CRTISO* (K), and *PhyB2* (L). AIC takes into account the likelihood of the models as well as its complexity, i.e. the number of parameters in the model, to establish which model is best supported by the data. Delta AIC values higher than 0 indicate higher support for the simpler model (3-state model without perfect repair), while negative values support the more complex model (3-state model). A Delta AIC of ~-6 corresponds to a relative likelihood of 0.05, indicating strong support for the more complex model (see related Figure 4, Table S2).

**Table S2.** Rate Constants estimated for kinetic model without precise repair.

| target | Rate | rate constant<sup>a</sup><br>(proportion events/hour) | CI<sup>a,c</sup><br>(1%) | CI<sup>a,c</sup><br>(99%) | flow after 72h<sup>b</sup><br>(proportion) | CI<sup>b,c</sup><br>(1%) | CI<sup>b,c</sup><br>(99%) |
| --- | --- | --- | --- | --- | --- | --- | --- |
| *Psy1* | Cutting | 0.0205 | 0.0117 | 0.0334 | 0.2192 | 0.1942 | 0.2331 |
|  | Repair-error | 0.0341 | 0.0278 | 0.0416 | 0.1581 | 0.1433 | 0.1705 |
|  | er1 | 0.2035 | 0.1631 | 0.2406 | NA | NA | NA |
|  | U (induction) | 0.7160 | 0.1902 | 0.7541 | NA | NA | NA |
|  | r (induction) | 27778.8235 | 0.0000 | 46083.6335 | NA | NA | NA |
|  | decay (induction) | 0.0000 | 0.0000 | 0.0000 | NA | NA | NA |
| *PhyB2* | Cutting | 0.0504 | 0.0422 | 0.0584 | 0.5453 | 0.5163 | 0.5805 |
|  | Repair-error | 0.0850 | 0.0731 | 0.1046 | 0.5250 | 0.4941 | 0.5598 |
|  | er1 | 0.0809 | 0.0469 | 0.1364 | NA | NA | NA |
|  | U (induction) | 0.4395 | 0.3944 | 0.4731 | NA | NA | NA |
|  | r (induction) | 32.0782 | 12.3336 | 61983.8014 | NA | NA | NA |
|  | decay (induction) | 0.0000 | 0.0000 | 0.0000 | NA | NA | NA |
| *CRTISO* | Cutting | 0.0171 | 0.0122 | 0.0254 | 0.5010 | 0.4055 | 0.5816 |
|  | Repair-error | 0.0803 | 0.0661 | 0.0976 | 0.4456 | 0.3633 | 0.5219 |
|  | er1 | 0.3131 | 0.2281 | 0.3824 | NA | NA | NA |
|  | U (induction) | 0.2914 | 0.0000 | 0.4850 | NA | NA | NA |
|  | r (induction) | 30886.4754 | 0.0000 | 74548.3365 | NA | NA | NA |
|  | decay (induction) | 0.0000 | 0.0000 | 0.0053 | NA | NA | NA |

Estimates for the rate constants and the three parameters of the induction curve, uncut fraction (u), induction speed (r), and induction decay (see Methods). <sup>a</sup>Rates are reported as the number of events per molecule per hour, with the exception of the rate of the steepness of the induction curve (r induction), which is reported as the number of events in 0.001 molecules. <sup>b</sup>The flow is reported as the proportion of molecules that experienced that event at the end of the experiment. <sup>c</sup>Confidence intervals (CI) are reported as the 1% and 99% percentiles of the estimates obtained from 100 stratified bootstraps of the data.

**Figure S4.**
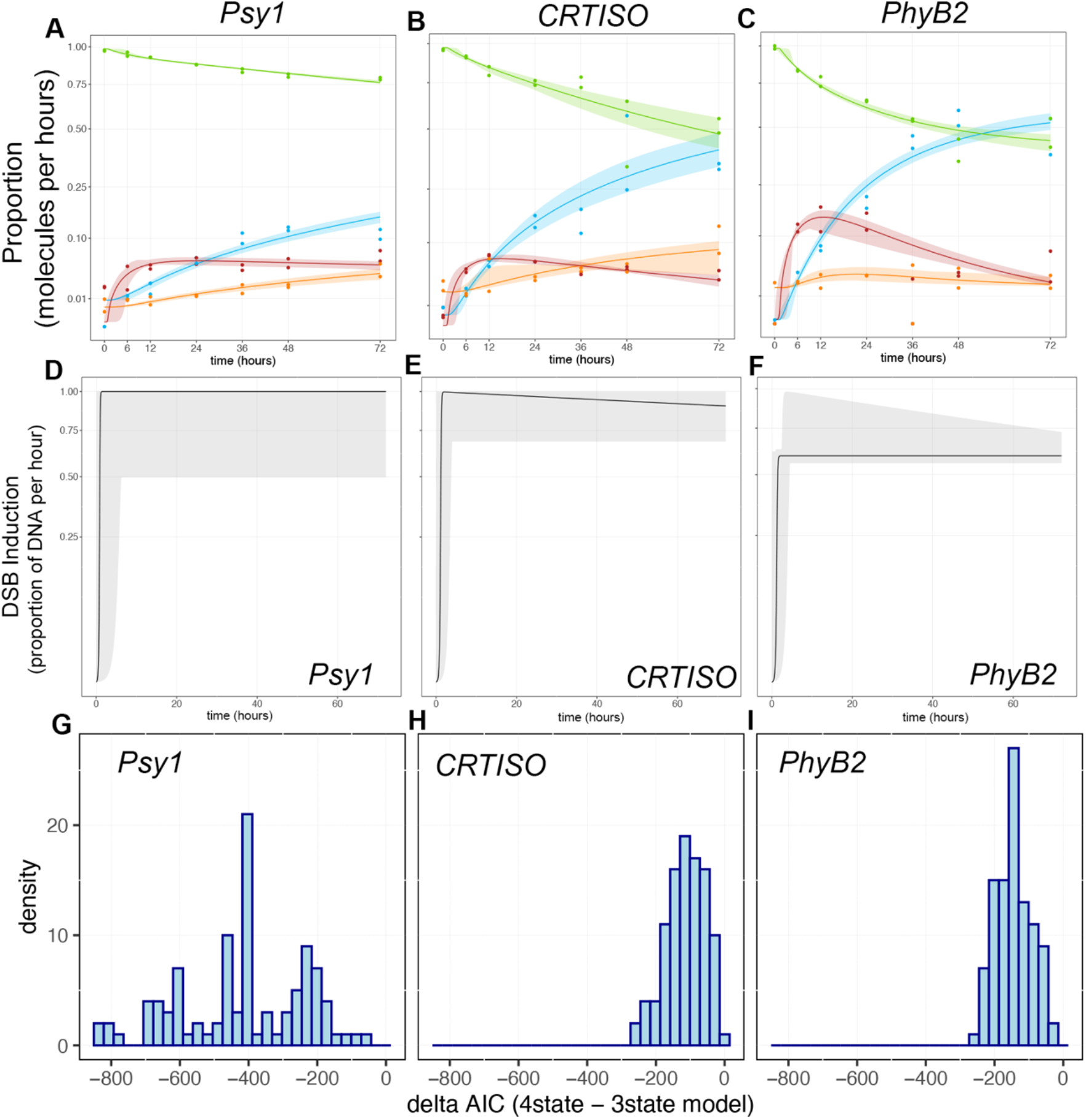
4-state model of DSB induction and repair. A-C) Fit of the model to the data at *Psy1* (A), *CRTISO* (B), and *PhyB2* (C). CIs represented in shading and calculated from 100 iterations of the boostrapping. D-F) Induction curves with confidence intervals from the bootstrap indicated in gray shading at *Psy1* (D), *CRTISO* (E), and *PhyB2* (F). G-I) Difference in AIC (Delta AIC, where AIC stands for Akaike Information Criterion) between the 4-state model and the 3-state model at *Psy1* (G), *CRTISO* (H), and *PhyB2* (I). AIC takes into account the likelihood of the models as well as its complexity, i.e. the number of parameters in the model, to establish which model is best supported by the data. Delta AIC values higher than 0 indicate higher support for the simpler model (3-state model), while negative values support the more complex model (4-state model). A Delta AIC of ~-6 corresponds to a relative likelihood of 0.05, indicating strong support for the more complex model (see related Figure 6, Table S3)

**Table S3.**
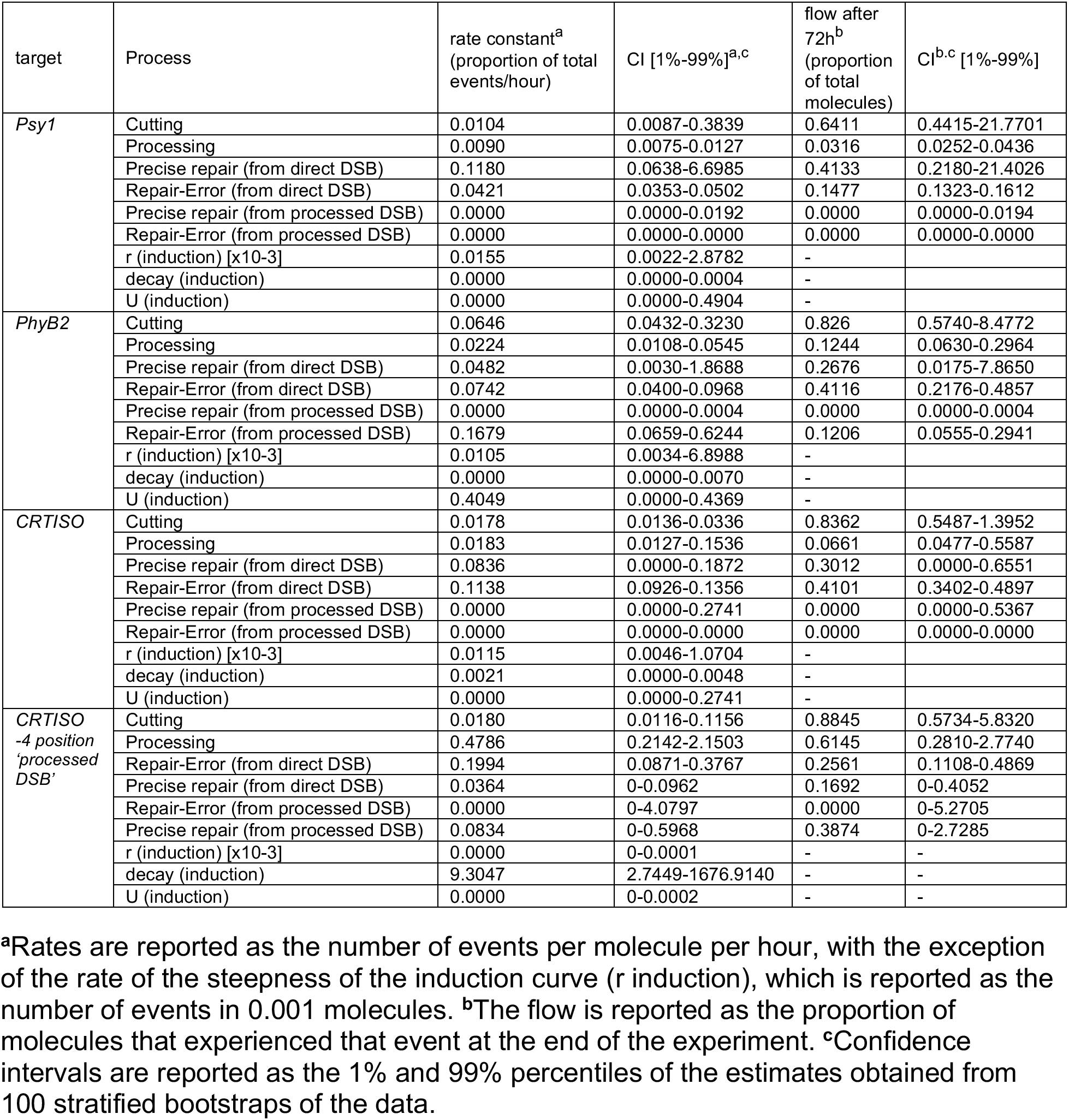
Rates and Flows Estimated by a 4-State Model of DSB Repair.

| target | Process | rate constant<sup>a</sup><br>(proportion of total events/hour) | CI [1%-99%]<sup>a,c</sup> | flow after 72h<sup>b</sup><br>(proportion of total molecules) | CI<sup>b,c</sup> [1%-99%] |
| --- | --- | --- | --- | --- | --- |
| *Psy1* | Cutting | 0.0104 | 0.0087-0.3839 | 0.6411 | 0.4415-21.7701 |
|  | Processing | 0.0090 | 0.0075-0.0127 | 0.0316 | 0.0252-0.0436 |
|  | Precise repair (from direct DSB) | 0.1180 | 0.0638-6.6985 | 0.4133 | 0.2180-21.4026 |
|  | Repair-Error (from direct DSB) | 0.0421 | 0.0353-0.0502 | 0.1477 | 0.1323-0.1612 |
|  | Precise repair (from processed DSB) | 0.0000 | 0.0000-0.0192 | 0.0000 | 0.0000-0.0194 |
|  | Repair-Error (from processed DSB) | 0.0000 | 0.0000-0.0000 | 0.0000 | 0.0000-0.0000 |
|  | r (induction) [x10-3] | 0.0155 | 0.0022-2.8782 | - | - |
|  | decay (induction) | 0.0000 | 0.0000-0.0004 | - | - |
|  | U (induction) | 0.0000 | 0.0000-0.4904 | - | - |
| *PhyB2* | Cutting | 0.0646 | 0.0432-0.3230 | 0.826 | 0.5740-8.4772 |
|  | Processing | 0.0224 | 0.0108-0.0545 | 0.1244 | 0.0630-0.2964 |
|  | Precise repair (from direct DSB) | 0.0482 | 0.0030-1.8688 | 0.2676 | 0.0175-7.8650 |
|  | Repair-Error (from direct DSB) | 0.0742 | 0.0400-0.0968 | 0.4116 | 0.2176-0.4857 |
|  | Precise repair (from processed DSB) | 0.0000 | 0.0000-0.0004 | 0.0000 | 0.0000-0.0004 |
|  | Repair-Error (from processed DSB) | 0.1679 | 0.0659-0.6244 | 0.1206 | 0.0555-0.2941 |
|  | r (induction) [x10-3] | 0.0105 | 0.0034-6.8988 | - | - |
|  | decay (induction) | 0.0000 | 0.0000-0.0070 | - | - |
|  | U (induction) | 0.4049 | 0.0000-0.4369 | - | - |
| *CRTISO* | Cutting | 0.0178 | 0.0136-0.0336 | 0.8362 | 0.5487-1.3952 |
|  | Processing | 0.0183 | 0.0127-0.1536 | 0.0661 | 0.0477-0.5587 |
|  | Precise repair (from direct DSB) | 0.0836 | 0.0000-0.1872 | 0.3012 | 0.0000-0.6551 |
|  | Repair-Error (from direct DSB) | 0.1138 | 0.0926-0.1356 | 0.4101 | 0.3402-0.4897 |
|  | Precise repair (from processed DSB) | 0.0000 | 0.0000-0.2741 | 0.0000 | 0.0000-0.5367 |
|  | Repair-Error (from processed DSB) | 0.0000 | 0.0000-0.0000 | 0.0000 | 0.0000-0.0000 |
|  | r (induction) [x10-3] | 0.0115 | 0.0046-1.0704 | - | - |
|  | decay (induction) | 0.0021 | 0.0000-0.0048 | - | - |
|  | U (induction) | 0.0000 | 0.0000-0.2741 | - | - |
| *CRTISO*<br>-4 position<br>'processed<br>DSB' | Cutting | 0.0180 | 0.0116-0.1156 | 0.8845 | 0.5734-5.8320 |
|  | Processing | 0.4786 | 0.2142-2.1503 | 0.6145 | 0.2810-2.7740 |
|  | Repair-Error (from direct DSB) | 0.1994 | 0.0871-0.3767 | 0.2561 | 0.1108-0.4869 |
|  | Precise repair (from direct DSB) | 0.0364 | 0-0.0962 | 0.1692 | 0-0.4052 |
|  | Repair-Error (from processed DSB) | 0.0000 | 0-4.0797 | 0.0000 | 0-5.2705 |
|  | Precise repair (from processed DSB) | 0.0834 | 0-0.5968 | 0.3874 | 0-2.7285 |
|  | r (induction) [x10-3] | 0.0000 | 0-0.0001 | - | - |
|  | decay (induction) | 9.3047 | 2.7449-1676.9140 | - | - |
|  | U (induction) | 0.0000 | 0-0.0002 | - | - |

<sup>a</sup>Rates are reported as the number of events per molecule per hour, with the exception of the rate of the steepness of the induction curve (r induction), which is reported as the number of events in 0.001 molecules. <sup>b</sup>The flow is reported as the proportion of molecules that experienced that event at the end of the experiment. <sup>c</sup>Confidence intervals are reported as the 1% and 99% percentiles of the estimates obtained from 100 stratified bootstraps of the data.

**Figures S5.**
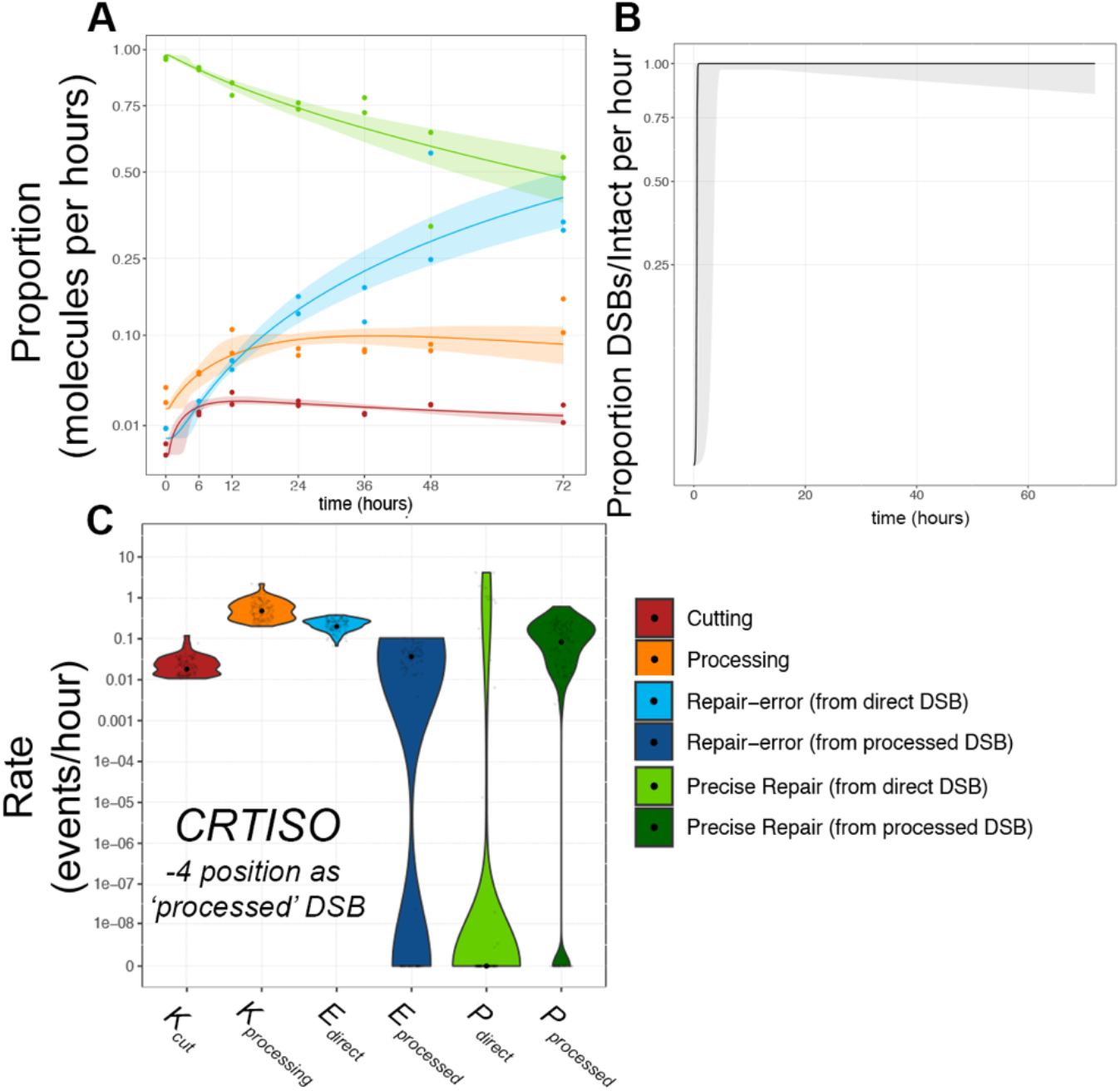
Complex Dynamics at CRTISO. A) Fit of the model to the data for CRTISO when the -4 bp positioned DSB is considered as ‘processed’ with CIs represented in shading and calculated from 100 iterations of the bootstrapping. B) Induction curve with CIs from boostrapping indicated in gray shading. C) Violin plots of rate constants estimated in terms of proportion per hour, smoothed distribution of the estimates obtained through the bootstrap procedure. Mean estimate of 100 iterations of bootstrapping is shown as a black point

Supplementary File 1. Table of oligonucleotides used in this study

Supplementary File 2. Excel workbook of consensus sequence counts for each dataset in the study

