## Supplementary File 1 for "Uncovering the Dynamics of Precise Repair at CRISPR/Cas9-induced Double-Strand Breaks"

**Supplementary File 1: Oligonucleotides used in this study**

| **Oligonucleotides** | | | | |
| --- | --- | --- | --- | --- |
| *UMI-DSBseq Adaptor sequences* | | | | |
| Adaptor Short Tail | | GATCGGAAGAGCGGGGACTATTTGC | IDT Ultramer DNA Oligos | |
| Adaptor long tail UMI barcode i7_1 | | CAAGCAGAAGACGGCATACGAGATNNNNNNNNNACGATCAGGTGACTGGAGTTCAGACGTGTGCTCTTCCGATC*T  (*denotes phosphorothioate bond) | IDT Ultramer DNA Oligos | |
| Adaptor long tail UMI barcode i7_2 | | CAAGCAGAAGACGGCATACGAGATNNNNNNNNNTCGAGAGTGTGACTGGAGTTCAGACGTGTGCTCTTCCGATC*T  (*denotes phosphorothioate bond) | IDT Ultramer DNA Oligos | |
| Adaptor long tail UMI barcode i7_3 | | CAAGCAGAAGACGGCATACGAGATNNNNNNNNNCTAGCTCAGTGACTGGAGTTCAGACGTGTGCTCTTCCGATC*T  (*denotes phosphorothioate bond) | IDT Ultramer DNA Oligos | |
| Adaptor long tail UMI barcode i7_4 | | CAAGCAGAAGACGGCATACGAGATNNNNNNNNNATCGTCTCGTGACTGGAGTTCAGACGTGTGCTCTTCCGATC*T  (*denotes phosphorothioate bond) | IDT Ultramer DNA Oligos | |
| Adaptor long tail UMI barcode i7_5 | | CAAGCAGAAGACGGCATACGAGATNNNNNNNNNTCGACAAGGTGACTGGAGTTCAGACGTGTGCTCTTCCGATC*T  (*denotes phosphorothioate bond) | IDT Ultramer DNA Oligos | |
| Adaptor long tail UMI barcode i7_6 | | CAAGCAGAAGACGGCATACGAGATNNNNNNNNNCCTTGGAAGTGACTGGAGTTCAGACGTGTGCTCTTCCGATC*T  (*denotes phosphorothioate bond) | IDT Ultramer DNA Oligos | |
| Adaptor long tail UMI barcode i7_7 | | CAAGCAGAAGACGGCATACGAGATNNNNNNNNNATCATGCGGTGACTGGAGTTCAGACGTGTGCTCTTCCGATC*T  (*denotes phosphorothioate bond) | IDT Ultramer DNA Oligos | |
| Adaptor long tail UMI barcode i7_8 | | CAAGCAGAAGACGGCATACGAGATNNNNNNNNNTGTTCCGTGTGACTGGAGTTCAGACGTGTGCTCTTCCGATC  (*denotes phosphorothioate bond) | IDT Ultramer DNA Oligos | |
| Adaptor long tail UMI barcode i7_9 | | CAAGCAGAAGACGGCATACGAGATNNNNNNNNNATTAGCCGGTGACTGGAGTTCAGACGTGTGCTCTTCCGATC*T  (*denotes phosphorothioate bond) | IDT Ultramer DNA Oligos | |
| Adaptor long tail UMI barcode i7_10 | | CAAGCAGAAGACGGCATACGAGATNNNNNNNNNCGATCGATGTGACTGGAGTTCAGACGTGTGCTCTTCCGATC*T  (*denotes phosphorothioate bond) | IDT Ultramer DNA Oligos | |
| Adaptor long tail UMI barcode i7_11 | | CAAGCAGAAGACGGCATACGAGATNNNNNNNNNGATCTTGCGTGACTGGAGTTCAGACGTGTGCTCTTCCGATC*T  (*denotes phosphorothioate bond) | IDT Ultramer DNA Oligos | |
| Adaptor long tail UMI barcode i7_12 | | CAAGCAGAAGACGGCATACGAGATNNNNNNNNNAGGATAGCGTGACTGGAGTTCAGACGTGTGCTCTTCCGATC*T  (*denotes phosphorothioate bond) | IDT Ultramer DNA Oligos | |
| Adaptor long tail UMI barcode i7_13 | | CAAGCAGAAGACGGCATACGAGATNNNNNNNNNGTAGCGTAGTGACTGGAGTTCAGACGTGTGCTCTTCCGATC*T  (*denotes phosphorothioate bond) | IDT Ultramer DNA Oligos | |
| Adaptor long tail UMI barcode i7_14 | | CAAGCAGAAGACGGCATACGAGATNNNNNNNNNAGAGTCCAGTGACTGGAGTTCAGACGTGTGCTCTTCCGATC*T  (*denotes phosphorothioate bond) | IDT Ultramer DNA Oligos | |
| **Primer Sequences** | | | | |
| P5 Target specific primer for PhyB2  P5 PhyB2 exon1 DSB2 Rev | | AATGATACGGCGACCACCGAGATCTACACTCTTTCCCTACACGACGCTCTTCCGATCTNNNGTGGGTGAGTCTCGGAGAAG | This paper | |
| P5 Target specific primer for Psy1  P5 Psy1-1 HTseq F | | AATGATACGGCGACCACCGAGATCTACACTCTTTCCCTACACGACGCTCTTCCGATCTNNNGGTTTGCCTGTCTGTGGTCT | Filler-Hayut et al., 2017 | |
| P5 Target specific primer for CRTISO  *P5 CRT_e4_DSB F* | | AATGATACGGCGACCACCGAGATCTACACTCTTTCCCTACACGACGCTCTTCCGATCTNNNGATGGTCCTGAGTGTTTGCC | This paper | |
| P7 amplification and enrichment primer | | CAAGCAGAAGACGGCATACGAGAT | Blecher-Gonenet al.,2013 | |
| *P5 enrichment primer with barcode* | | AATGATACGGCGACCACCGAGATCTACAC**NNNNNNNN**ACACTCTTTCCCTACACGAC | Adapted from Blecher-Gonenet al.,2013 | |
| **crRNA** | | | | |
| *CRTISO* Alt-R® CRISPR-Cas9 crRNA | | GCGATGCTACCAGCATTCTG | Dahan-Meir et al., 2018 | |
| *Psy1* Alt-R® CRISPR-Cas9 crRNA | | GAATGTCTGTTGCCTTGTTA | Filler-Hayut et al., 2017 | |
| *PhyB2* Alt-R® CRISPR-Cas9 crRNA | | GGCCTGCATAAGGAATTCAC | This paper | |
| **Target Sequences** | | | | |
| ***Target*** | ***Amplicon*** | | | ***Restriction enzyme*** |
| *CRTISO* | TATTTAAGTGAAGAATGGTAAAGGGGAGGGGCCATTATCCCCGAGTTTTGAGCACTATTGATGGTCCTGAGTGTTTGCCCTCGGTCATCAAAAAAATTTAAGGAGTCATCTTTCACGCTGATGTGTGCAGCGCGCGACGTGCTTAATTATCCTACCGTAGAATCTTAATTTATGCCATCATTATTCATTACAGCCTACTATTTGCCCCAGAATGCTGGTAGCATCGCTCGGAAGTATATAAGAGATCCTGGGTTGCTGTCTTTTATAGATGCAGAG | | | MspI |
| *Psy1* | GGTTTGCCTGTCTGTGGTCTTTTTATAATCTTTTTCTACAGAAGAGAAAGTGGGTAATTTTGTTTGAGAGTGGAAATATTCTCTAGTGGGAATCTACTAGGAGTAATTTATTTTCTATAAACTAAGTAAAGTTTGGAAGGTGACAAAAAGAAAGACAAAAATCTTGGAATTGTTTTAGACAACCAAGGTTTTCTTGCTCAGAATGTCTGTTGCCTTGTTATGGGTTGTTTCTCCTTGTGACGTCTCAAATGGGACAAGTTTCATGGAATCAGTCCG | | | EcoRI-HF |
| *PhyB2* | GTGGGTGAGTCTCGGAGAAGCATGTCACATAACACTGTCTGTGTCCTCAAAACCCGCTTTTCAGCCAACTGCGATGCCAATTGTAATTCCATGTTTAGTTGGAGCCCAAAGGCCTGCATAAGGAATTCACAGGCATAACGAAGGGGGAAAGGAATGAACCGAGCTGAACTGTGGTGCCCAACAACCAAACCCCATAATCTCATTGCATTTCGGCCTCCAACAACTTCATCATCATTTCCATTTA | | | AanI-FD |
